# Structure-guided microbial targeting of antistaphylococcal prodrugs

**DOI:** 10.1101/2020.12.15.408237

**Authors:** Justin J. Miller, Ishaan T. Shah, Jayda Hatten, Yasaman Barekatain, Elizabeth A. Mueller, Ahmed M. Moustafa, Rachel L. Edwards, Cynthia S. Dowd, Paul J. Planet, Florian L. Muller, Joseph M. Jez, Audrey R. Odom John

## Abstract

Carboxy ester prodrugs have been widely employed as a means to increase oral absorption and potency of phosphonate antibiotics. Prodrugging can successfully mask problematic chemical features that prevent cellular uptake and can be used to target delivery of compounds to specific tissues. However, many carboxy ester promoieties are rapidly hydrolyzed by serum esterases, curbing their potential therapeutic applications. While carboxy ester-based prodrug targeting is feasible, it has seen limited use in microbes due to a paucity of information about the selectivity of microbial esterases. Here we identify the bacterial esterases, GloB and FrmB, that are required for carboxy ester prodrug activation in *Staphylococcus aureus.* Additionally, we determine the substrate specificities for FrmB and GloB and demonstrate the structural basis of these preferences. Finally, we establish the carboxy ester substrate specificities of human and mouse sera, which revealed several promoieties likely to be serum esterase-resistant while still being microbially labile. These studies lay the groundwork for structure-guided design of anti-staphyloccal promoieties and expand the range of molecules to target staphyloccal pathogens.

## Introduction

Antimicrobial resistance presents a major challenge to public health^1,2^. In 2019, 2.8 million antibiotic-resistant infections occurred in the United States, resulting in 35,000 deaths^3^. Some estimates have suggested that antimicrobial-resistant infections will cause as many as 10 million deaths annually by 2050^4^. *Staphylococcus aureus* is a devastating human pathogen. Methicillin-resistant *S. aureus* (MRSA) has been labeled a “serious threat” by the Centers for Disease Control and Prevention^3,5,6^. New antimicrobials, especially those with novel mechanisms of action, are urgently needed; however, most anti-infectives under development take advantage of existing antibiotic scaffolds with proven efficacy and established safety profiles^7,8^. New, chemically distinct antibiotics are highly desirable as a strategy to circumvent antimicrobial resistance.

Many metabolic processes are essential for microbial growth and/or pathogenesis, although few existing antimicrobials exploit metabolism as an anti-infective target. Metabolic drug design can be facile, as natural substrates serve as a template for competitive inhibitors. As metabolism often involves the transformation of highly charged and polar metabolites, inhibitors of metabolic enzymes often take advantage of phosphonate functional groups for target engagement^9^. Unfortunately, negatively charged phosphonates are readily excluded from cell membranes and often exhibit poor drug-like properties^10–19^. New strategies enabling effective deployment of antimetabolites will serve to expand the druggable space for antimicrobials.

One means of improving phosphonate cellular permeability is to chemically mask the undesirable negative charge with lipophilic groups. This strategy (termed prodrugging) can employ labile promoieties that are removed during absorption, distribution, or intracellularly to yield the original phosphonate antibiotic (Figure 1a)^17–19^. We have previously designed a series of MEPicide antibiotics, in which phosphonate isoprenoid biosynthesis inhibitors are modified with a lipophilic pivaloyloxymethyl (POM) promoiety. MEPicide prodrugs bypass the need for active cellular transport, while simultaneously increasing compound potency against zoonotic staphylococci, *S. schleiferi* and *S. pseudintermedius*, as well as *Plasmodium falciparum* and *Mycobacterium tuberculosis*^11,13,20–23^. However, POM promoieties are rapidly hydrolyzed by serum carboxylesterases, which limits the *in vivo* efficacy of POM prodrugs as a strategy to improve the drug-like features of charged antimicrobials^12,24^.

**Figure 1.**
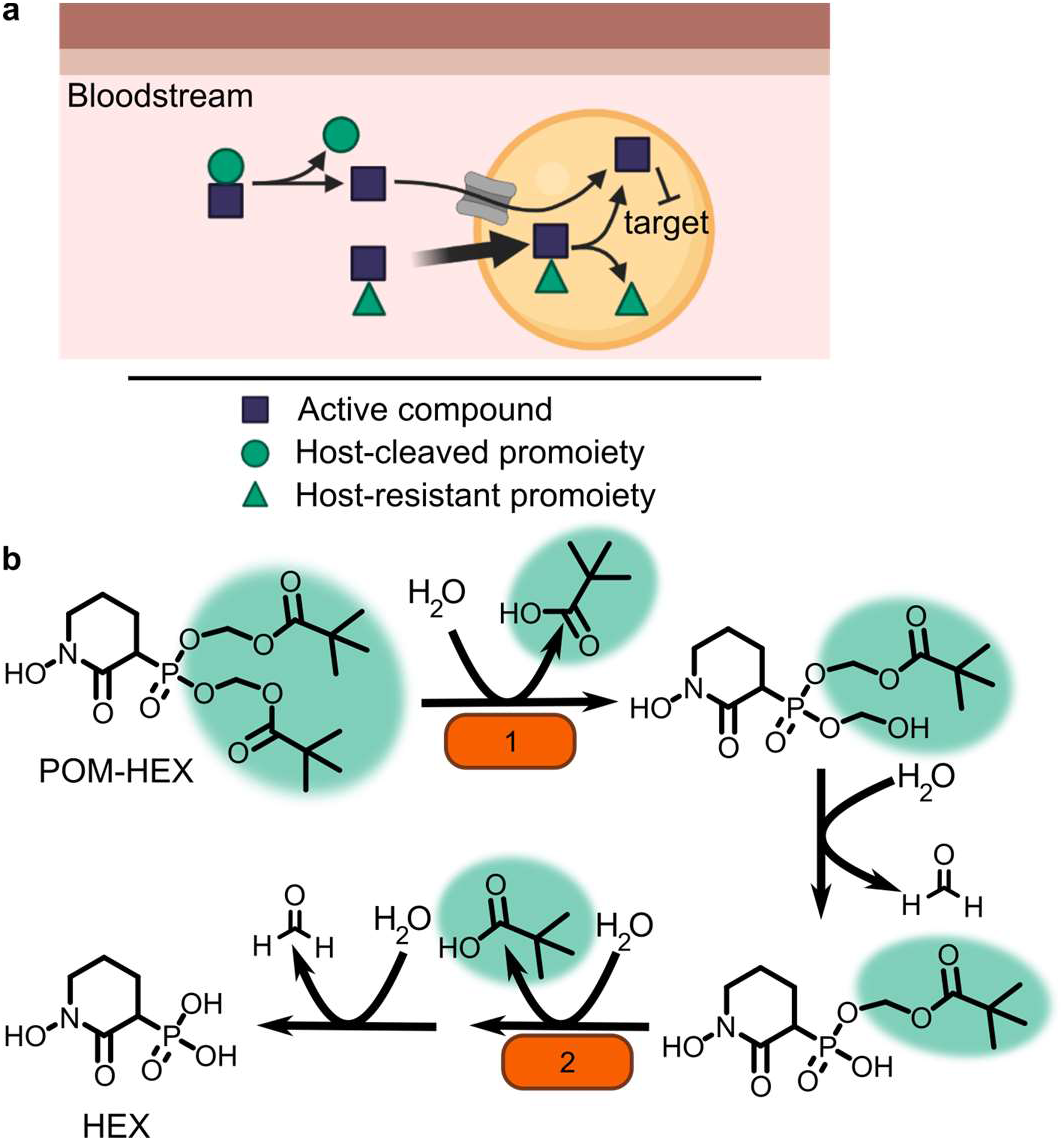
Prodrug activation model (a) and proposed enzymatic mechanism. (b) Carboxy ester promoieties highlighted in green.

To enable effective cellular delivery of phosphonate antibiotics, new lipophilic prodrug moieties that are resistant to serum carboxylesterases—yet cleavable by microbial esterases—are required. This strategy has been effectively used to achieve tissue-specific selective prodrug activation. For example, liver-targeted prodrug delivery was achieved by harnessing the distinct substrate specificity of the liver-specific isoform of P450, CYP3A4^25,26^. Accordingly, understanding the enzymatic and structural mechanisms of microbial prodrug activation will facilitate the development of microbe-specific prodrugs.

We recently described the staphylococcal enzyme, GloB, which facilitates activation of carboxy ester prodrugs in *S. schleiferi* and *S. pseudintermedius*^27^. Notably, GloB is insufficient to fully activate prodrugs *in vitro*, suggesting that at least one additional enzyme is necessary for complete phosphonate release. In this work, we identify and characterize the role of two *S. aureus* esterases, GloB and FrmB, which each catalyze carboxy ester prodrug hydrolysis and contribute to carboxy ester prodrug activation. We demonstrate that both esterases have defined substrate specificities, which diverge from the substrate specificities of human and mouse sera. Finally, we present the three-dimensional structures of GloB and FrmB to enable ongoing structure-guided design of FrmB- and GloB-targeted prodrugs.

## Results

### Identification of microbial esterases responsible for carboxylesterase activity

In the zoonotic staphylococcal species *S. schleiferi* and *S. pseudintermedius*, loss of the enzyme GloB, a hydroxyacylglutathione hydrolase, or glyoxalase II enzyme, confers resistance to carboxy ester prodrugs because carboxy ester prodrugs are not activated^27^. However, purified GloB alone is insufficient to activate carboxy ester prodrugs *in vitro*, suggesting that at least one additional cellular enzyme is required. Based on the hypothesized carboxy ester activation pathway, we predicted that the missing enzyme(s) might be either another carboxylesterase or a phosphodiesterase (Figure 1b). To establish the pathway for carboxy ester prodrug activation in *S. aureus*, we made use of the Nebraska Transposon Mutant Library (NTML), in which nearly 2,000 non-essential *S. aureus* genes have been individually disrupted by a stable transposon insertion^28^. Using the gene ontology feature on the NTML website (https://app1.unmc.edu/fgx/gene-ontologies.html), we identified six carboxylic ester hydrolases (including *gloB*), 11 phosphatases, and nine phosphoric diester hydrolases as candidate activators of carboxy ester prodrugs (Supplementary Table 1). Each identified transposon mutant was screened for resistance to the carboxy ester prodrug, POM-HEX. POM-HEX is a pivaloyloxymethyl prodrug of the compound HEX, which inhibits the glycolytic enzyme enolase (Figure 1b, Figure 2a). Of the 26 candidate esterase transposon mutants, only two strains were significantly more resistant to POM-HEX than the *S. aureus* parental strain, JE2, as determined by half-maximal growth inhibitory concentration (IC_50_) (Figure 2b, Supplementary Table 1). One of these strains had a transposon disrupting the gene that encodes the glyoxalase II enzyme, GloB. The GloB ortholog in *S. scheiferi* is a prodrug-activating enzyme, mutation of which confers resistance to POM-HEX^27^. The second POM-HEX-resistant strain harbored a transposon insertion in the locus encoding the predicted carboxylesterase annotated as FrmB. FrmB has been previously identified as FphF, a serine hydrolase, and is the primary *S. aureus* target of the sulfonyl fluoride compound JCP678^29^. As *S*-formylglutathione hydrolase activity is more likely the biological function of this protein, we will refer to this protein as it has been named in *E. coli* - FrmB.

**Figure 2.**
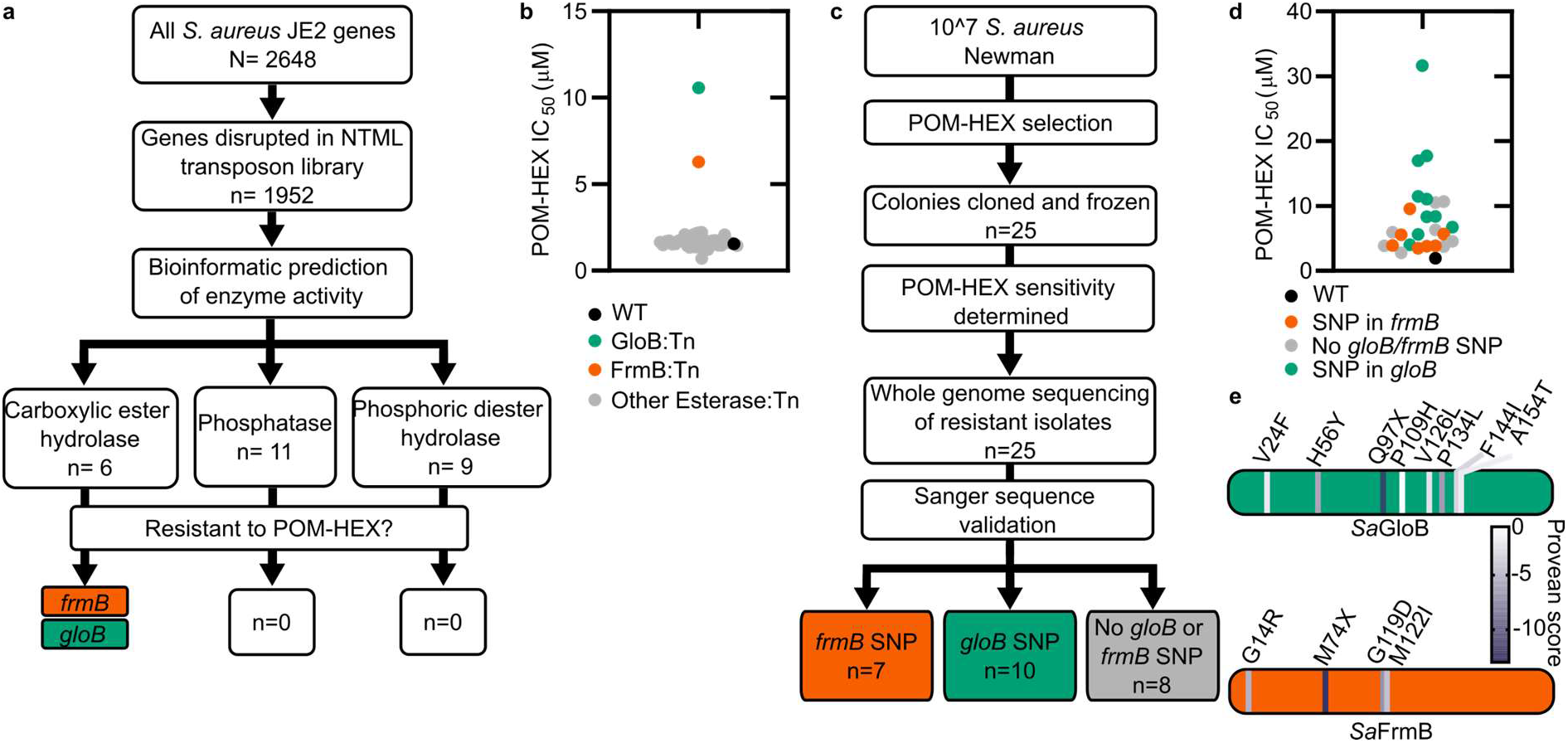
Forward and reverse genetics approaches identify FrmB and GloB as candidate POM-prodrug hydrolases in *S. aureus*. (a) Reverse genetics identification of candidate prodrug activating enzymes. (b) POM-HEX susceptibility of identified candidate resistance genes from (a) as determined by IC_50_. (c) Forward genetic screen approach. (d) POM-HEX susceptibility of POM-HEX-resistant *S. aureus*. (e) Nonsynonymous point mutations identified by whole-genome sequencing in *frmB* and *gloB*. In all experiments GloB is colored green and FrmB orange. Displayed are the means of three independent biological experiments.

In parallel, we employed an unbiased forward genetics approach to identify genetic changes associated with POM-HEX resistance. POM-HEX-resistant staphylococci were derived by exposing wild-type (WT) *S. aureus* Newman to growth inhibitory concentrations (IC) of POM-HEX. In total, we selected and cloned 25 isolates with IC_50_ values ranging from 1.5-16x that of WT *S. aureus* Newman (Figure 2 c, d, Supplementary Table 2). Whole genome sequencing of POM-HEX-resistant strains revealed mutations in *frmB* (n = 7), *gloB* (n = 10), and strains with no mutations in either *frmB* or *gloB* (n =8). Most mutations were nonsynonymous single nucleotide polymorphisms (SNPs) (Figure 2d, Supplementary Table 2). In three instances, *gloB* was the only verified genetic change in the genome. Additionally, *frmB* and *gloB* each had one instance of a mutation resulting in a premature stop codon truncating the protein to less than a 100 amino acid sequence. Strains with no identified mutations in either *frmB* or *gloB* did not have any mutations which are immediately obvious as POM-HEX resistance mechanisms. Of the non-*frmB* and non-*gloB* mutations, we selected transposon mutants from the NTML that had transposon insertions in the corresponding genes. These transposon mutations did not confer resistance to POM-HEX (Supplementary Table 3). Overwhelmingly, the observed mutations in both *frmB* and *gloB* are predicted to have deleterious effects on protein function (PROVEAN score below a threshold of −2.5) (Figure 2e).

To evaluate the sequence conservation of FrmB and GloB among *S. aureus*, we performed a WhatsGNU analysis on all publicly available *S. aureus* genomes. WhatsGNU is a bioinformatic tool that can compress large databases and assess the number of instances in which a specific gene has 100% sequence coverage and identity match within the entire database^30^. This parameter, the gene novelty unit (GNU) score, is high when a sequence is under strong selective pressure within the population, and low when the gene is variable. GloB exhibits an exceptionally high GNU score of 8215 (of 10350 possible) indicating that there is strong sequence conservation for *S*. *aureus* GloB. Conversely, FrmB sequences appear to be extremely conserved within individual *S. aureus* clonal complexes but varied between each complex (GNU scores of 2218 or 3370 of 10350, Supplementary Figure 1). We constructed a phylogenetic tree of GloB and FrmB orthologs among microbial populations. Most bacterial species contain a GloB ortholog, though the primary sequence is highly variable across species and does not readily cluster according to the tree of life (Supplementary Figure 2). FrmB sequences are also highly sequence divergent, though they tend to cluster closer to the expected tree of life (Supplementary Figure 2).

The agreement between our forward and reverse genetic screens strongly suggests that two discrete predicted esterases and not a pool of redundant cellular esterases activates the prodrug in *S. aureus*. Additionally, the finding that mutation in either *frmB* or *gloB* is sufficient to confer POM-HEX resistance suggests that the two enzymes may work in concert to bioconvert POM-HEX into HEX. As substantial sequence variation occurs among *frmB* and *gloB*, any prodrug therapeutics which hijack these two enzymes may not be broadly activated across all bacteria.

### FrmB and GloB are carboxylesterases with diverging substrate specificity

GloB is predicted to be a type II glyoxalase and a member of the large metallo-β-lactamase protein superfamily (INTERPRO IPR001279). Glyoxalase II enzymes, including the closely related GloB ortholog from *S. scheiferi*, catalyze the second step in the glyoxalase pathway that is responsible for the cellular conversion of methylglyoxal (a toxic glycolytic byproduct) to lactic acid^27,31–33^. Conversely, FrmB orthologs hydrolyze p-nitrophenyl esters of short-chain fatty acids (C2-C6) and are thought to mediate detoxification of cellular formaldehyde^34,35^.

We purified recombinant WT *Sa*FrmB and *Sa*GloB to evaluate the enzymatic function of each protein (Supplementary Figure 3a). We first assessed glyoxalase II activity using an assay, which couples hydrolysis of the glyoxalase II substrate, *S*-lactoylglutathione, to a change in absorbance (Supplementary Figure 3b). *Sa*GloB hydrolyzes *S*-lactoylglutathione with a specific activity comparable to previously characterized microbial type II glyoxalases, but *Sa*FrmB lacks appreciable glyoxalase activity.

We next assessed the ability of FrmB and GloB to hydrolyze p-nitrophenyl esters of short-chain fatty acids that have a photometric change upon hydrolysis (Supplementary Figure 3c). FrmB has modest activity against 4-nitrophenyl acetate and butyrate but no activity against 4-nitrophenyl trimethylacetate, suggesting a preference for unbranched fatty acids (Supplementary Figure 3c). This finding is in agreement with a previous characterization of FrmB as having a preference for short-chain and unbranched hydrophobic lipid substrates^35^. Notably, GloB has no detectable activity against these substrates and neither GloB nor FrmB hydrolyzes 4-nitrophenyl trimethylacetate despite its structural similarity to POM-HEX. This may be due to the absence of the acyloxymethyl ether moiety in 4-nitrophenyl substrates, which is found in POM-prodrugs.

We also sought to directly assess the role of GloB and FrmB in POM-HEX activation. We incubated each enzyme with POM-HEX and characterized the products via ^31^P-^1^H-heteronuclear single quantum coherence (HSQC) nuclear magnetic resonance (NMR). We have previously shown that GloB removes only one POM moiety, resulting in an accumulation of mono-POM-HEX (Figure 1b)^27^. Similarly, FrmB is capable of removing only one POM-moiety (Supplementary Figure 4). We hypothesized that the two esterases may be stereoselective and incubated both enzymes with POM-HEX. We find that incubation of POM-HEX with GloB and FrmB still results in an accumulation of mono-POM-HEX, suggesting that the two esterases may be unable to cleave the charged mono-POM species (Supplementary Figure 4).

### GloB and FrmB substrate specificity

To facilitate design of microbially targeted prodrug activation using these two enzymes, we next sought to extensively characterize GloB and FrmB substrate specificity. We employed a 32-compound ester substrate library, which fluoresces upon esterase activity (Supplementary Figure 5)^36^. This library systematically varies ester substrate length, branching patterns, and ether and sulfide positioning, thereby allowing for the precise determination of structure-activity relationships. Kinetic measurements were performed for both FrmB and GloB over a range of substrate concentrations for the entire library, allowing for the calculation of catalytic efficiency (*k*_cat_/*K*_m_) for each enzyme and substrate (Supplementary tables S4, S5).

We find that FrmB and GloB tend to have the highest activity toward oxygen ethers (Supplementary Figures 5 and 6). GloB has the highest activity against short-chain ethers (compounds 1-3), with some tolerance for branching at the first carbon beyond the ester carbonyl (compounds 7-9), while extensive branching strongly reduces activity (compound 10). Remarkably, GloB is tolerant of the extreme steric bulk introduced with the phenoxyacetic acid substrate, if the substrate contains an oxygen or sulfur ether (compound series 11). GloB exhibits a strong preference for oxygen at the β-position to the carbonyl over the γ-position but is indifferent to the positioning of sulfur. While GloB has a wider range of catalytic specificities, FrmB exhibits lower overall activity and a narrower range of catalytic efficiency. FrmB hydrolyzes unbranched substrates with little regard for chain length or the end-of-chain bulk within the tested substrates (compound series 1-3, 11). Branching at the position following the ester carbonyl (compound series 7-9, 12) is deleterious to FrmB activity. When oxygen is included in the chain, positioning at the β-position to the carbonyl is strongly preferred over the γ-position. For both GloB and FrmB, expanding the library of substrates tested will be revelatory of the true limitations of enzymatic activity.

### Importance of substrate specificity *in vivo*

While *in vitro* enzymatic substrate profiling is informative for how individual enzymes activate prodrugs, it may not reflect the complex biochemical processes happening *in vivo*, where additional cellular esterases may impact overall compound activation. We performed quantitative live cell imaging to measure the activation of pro-fluorescent substrates in real time. We loaded *S. aureus* onto a microfluidic device and imaged the cells by phase contrast and epifluorescence microscopy before and while supplying pro-fluorescent compound. Intracellular fluorescence accumulates in response to the rapid introduction of substrate into the chamber and can be quantified through time.

We selected four pro-fluorescent substrates of varying catalytic efficiency against FrmB and GloB to test in our microfluidics experiments. *In vitro*, *s*ubstrate 1O displays high activity for both FrmB and GloB, 3C displays moderate activity for FrmB and GloB, 5O has moderate activity against GloB but poor activity against FrmB, and 9C has poor activity against both GloB and FrmB (Supplementary Figures 5 and 6). Comparing the activation of these substrates through time, we find that our *in vitro* catalytic efficiency determination correlates well with our intracellular activation rates (Figure 3, Supplementary Movies 1-4). Compound 1O, which exhibits high catalytic efficiency with both GloB and FrmB, reaches fluorescence saturation before the first 2-minute time point. Compound 3C, which GloB and FrmB use with moderate activity, slowly activates over the duration of the experiment, and 5O and 9C, which have moderate-to-poor turnover with both GloB and FrmB, never appreciably activates during the 30 min of observation (Figure 3). As fluorescent activation is quantified per individual cell, we can also assess the uniformity of prodrug activation across the population. We observe remarkably homogenous activation of prodrugs across all observed cells (Figure 3).

**Figure 3.**
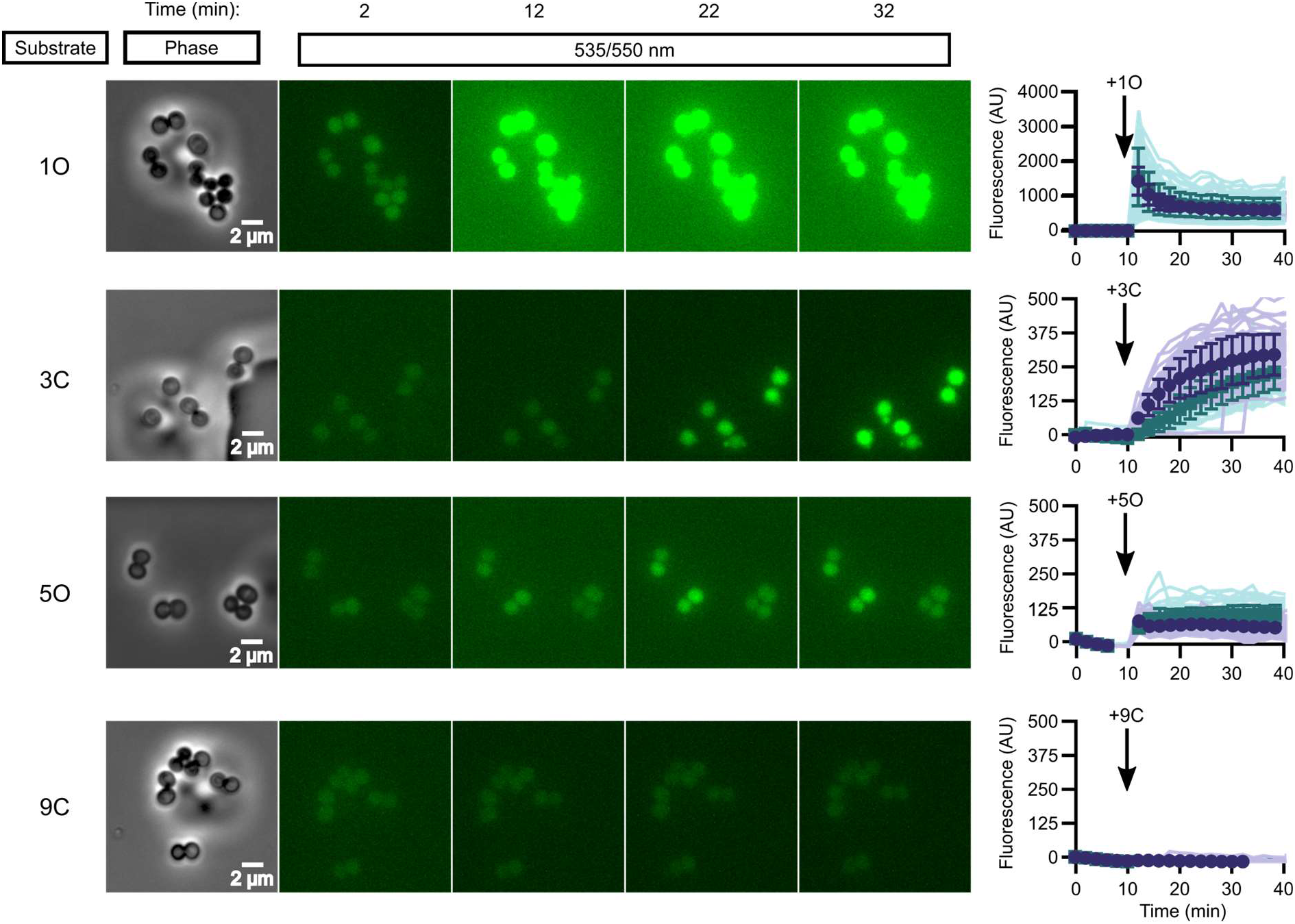
*In vivo* activation rates depend on ester promoiety selection. Time series of activation of various fluorogenic substrates (Supplementary Figure 5). Substrates were added into the microfluidics chamber at t = 10 minutes. On the right, quantification of individual cell or cell cluster fluorescence per area. Faint traces are individual cells and darker traces represent the mean of a given experiment. Each experiment was performed in biological duplicate and each experiment is displayed in a different color (purple or green). Error bars denote SD.

### Three-dimensional structure of FrmB

To establish the structural basis for FrmB and GloB substrate specificity and enable future structure-guided prodrug therapeutic design, we solved the three-dimensional structures of both *i>S. aureus* FrmB and GloB. *S. aureus* FrmB was solved at 1.60 Å using molecular replacement with the low-temperature active alkaline esterase Est12 (PDB ID: 4RGY) as a search model^37^. Refinement parameters and statistics are summarized in Supplementary Table 6. A single dimer of FrmB is observable in the asymmetric unit, matching the apparent molecular weight of FrmB as observed via size-exclusion chromatography. The overall fold of FrmB is characteristic of the α/β-hydrolase fold. Six parallel β-strands and one anti-parallel β-strand pair form a central eight stranded β-sheet, which is surrounded by α-helices (Figure 4a). One monomer of FrmB has electron density for a single magnesium ion, whereas the second monomer has two magnesium ions present.

**Figure 4.**
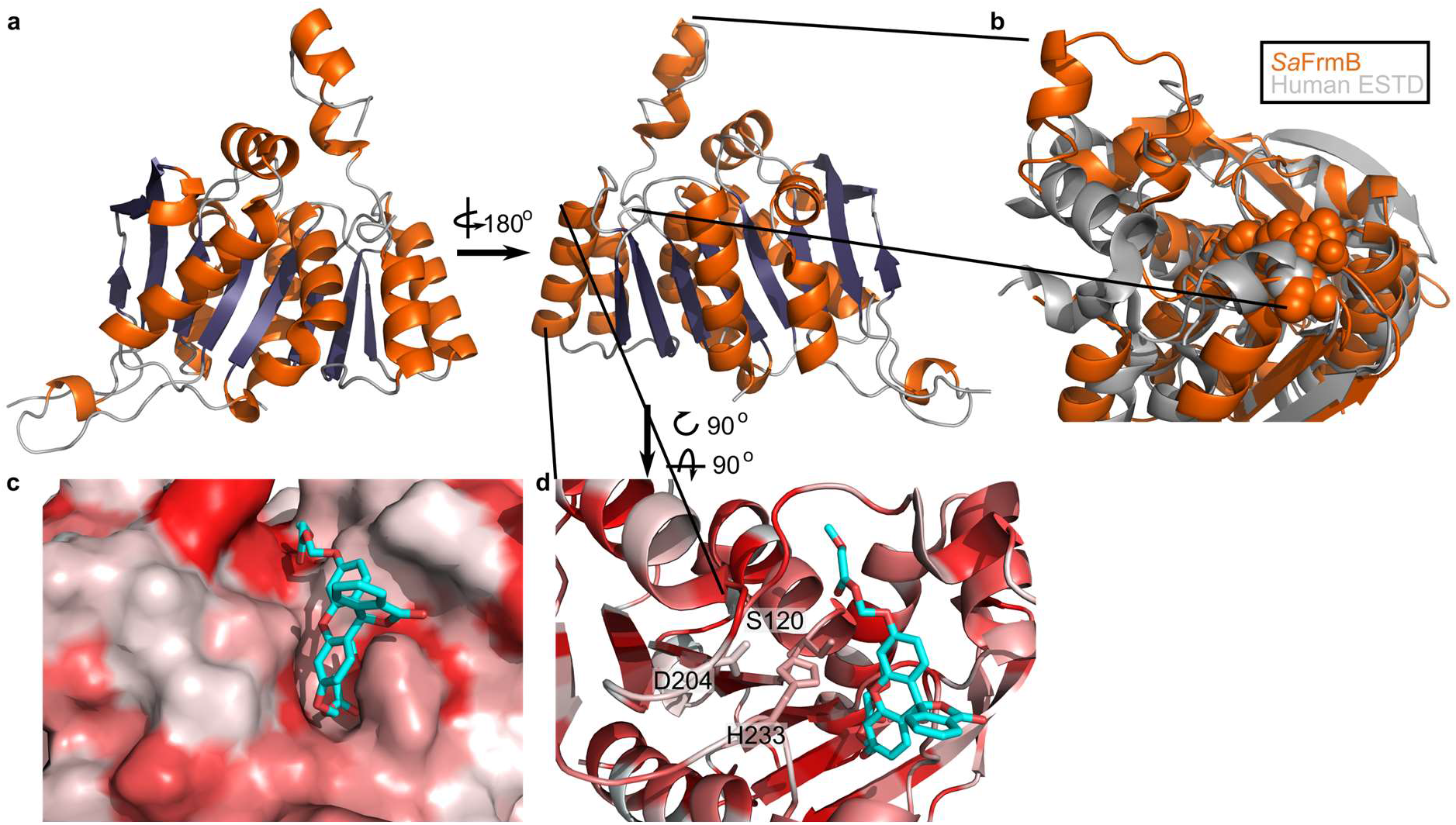
Three-dimensional structure of FrmB. (a) Overall fold, α-helices colored in orange and β-strands colored in purple. (b) Comparison between *Sa*FrmB (orange) and its closest human ortholog, ESTD (gray). Active site residues denoted in orange spheres. (c, d) Docking of substrate 1O (sticks) in the active site of FrmB. surface view, red indicates highly hydrophobic and white hydrophilic residues. Surface view (c) or stick view with catalytic triad (d).

A structural similarity search was performed using the DALI server to identify proteins related to *Sa*FrmB. The structure of *Sa*FrmB was most similar to the molecular replacement model, Est12 from deep sea bacteria (PDB ID 4RGY, root mean squared deviation (r.m.s.d.) = 1.020 Å), but also had similarity to *Bacteroides intestinalis* ferulic acid esterase *Bi*Fae1A (PDB ID 5VOL,r.m.s.d. = 1.137 Å) and *Streptococcus pneumonia* tributyrin esterase estA (PDB ID 2UZ0, r.m.s.d. = 1.329 Å)^38–40^. All structures display strong structural conservation including the positioning of the prototypic serine hydrolase catalytic triad: Ser120, Asp204, and His233 (*S. aureus*) (Supplementary Figure 7). The most striking difference between the related structures is the flexible cap domain, implicated in substrate specificity of EstA and Est12^37,40^. While this manuscript was in preparation, an independent structure of FrmB was reported^35^. The two structures are nearly identical (PDB ID 6ZHD, r.m.s.d = 0.433 Å), with slightly different positioning of the capping domain.

We compared *Sa*FrmB to its closest human ortholog, human esterase D (PDB ID 3fcx), finding moderate structural similarity both in the overall fold (r.m.s.d = 4.625 Å) and in the positioning of the catalytic triad^41^. However, *Sa*FrmB and human esterase D differ notably in the solvent-accessible surface around the active site, suggesting the potential for distinct substrate utilization, primarily driven by differential positioning of the cap domain (Figure 4b).

Using Autodock Vina, we modeled the highest catalytic efficiency substrate of FrmB, 1O, onto the active site of FrmB^42,43^. Serine hydrolases classically bind the substrate carbonyl oxygen in an oxyanion hole, and substrate hydrolysis is initiated through attack of the catalytic serine on the ester carbonyl. The docking of 1O on FrmB mimics the initial state of a serine hydrolase reaction, with the carbonyl oxygen buried and the catalytic serine poised for attack (Figure 4c). The pocket directly next to the oxyanion hole is relatively narrow, suggesting that steric hindrance explains the poor activity of FrmB against branched substrates. The active site pocket extends and opens significantly after passing by the oxyanion hole, supporting the capacity of FrmB to hydrolyze substrates that contain large steric groups distant from the carbonyl carbon, such as 11O.

### Three-dimensional structure of GloB

We also solved the structure of *Sa*GloB to 1.65 Å, using selenomethionine (SeMet)-substituted GloB and molecular replacement using metallo-β-lactamase TTHA1623 from *Thermus thermophilus* as a search model^44^. Final structural refinement parameters and statistics are summarized in Supplementary Table 6. Four monomers of *Sa*GloB are observed in the asymmetric unit with each displaying crystallographic symmetry. *Sa*GloB exhibits the classic αβ/βα-fold that defines the metallo-β-lactamases, including glyoxalase II (Figure 5a)^32,44–46^.

As with *Sa*FrmB, a DALI server search was performed to identify proteins structurally similar to *Sa*GloB. *Sa*GloB displays extremely high similarity to the unusual type II glyoxalase YcbI from *Salmonella enterica* (PDB ID: 2XF4, r.m.s.d = 0.898 Å), the molecular replacement search model TTHA1623 from *Thermus thermophilus* (PDB ID: 2ZWR, r.m.s.d. = 0.767), and to the *Arabidopsis thaliana* glyoxalase II (PDB ID: 1XM8, r.m.s.d = 1.165 Å), with the exception that *At*GloB has a 50 amino acid C-terminal extension (Supplementary Figure 8a)^32,44,47^. Also consistent with previously observed GloB structures, *Sa*GloB shows clear electron density for two zinc ions coordinated by six histidines and two aspartates (Supplementary Figure 8b). Density for a water molecule is also visible and appears to be coordinated by the two zinc ions, as observed for human glyoxalase II^45^.

Overlaying *S. aureus* GloB with *Homo sapiens* GloB (PDB: 1qh5) reveals that the two structures are remarkably similar (r.m.s.d = 1.249 Å), with a few notable exceptions. *Hs*GloB has two extensions: one, a 34-amino acid insertion; the other, a 32-amino acid C-terminal extension, both of which form α-helical features that abut the active site (Figure 5b)^45^. On the opposite side of the active site, *Sa*GloB has a 19-amino acid flexible loop which is partially observed in the electron density. This loop is positioned such that it may cover the active site to sterically hinder substrate access (Figure 5b). Overall, these structural differences between *Hs*GloB and *Sa*GloB support the design of prodrug substrates selectively cleaved by *S. aureus*.

**Figure 5.**
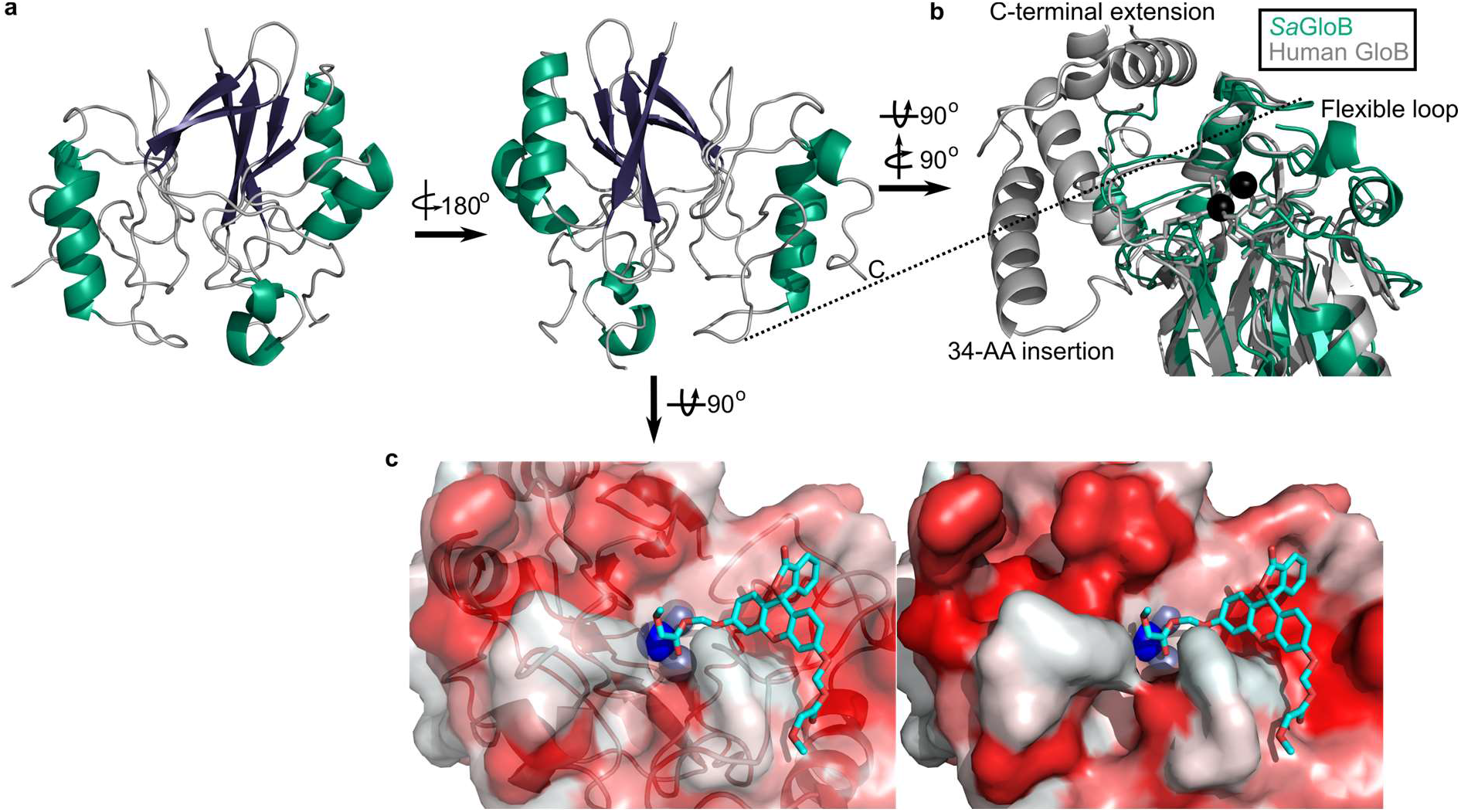
Three-dimensional structure of GloB. (a) Overall fold, α-alpha helices colored in green and β-strands colored in purple. (b) Comparison of *Sa*GloB (green) and human GloB (gray). (c) Docking of the substrate 1O (sticks) in the active site of GloB. Left, partial cartoon view, Right surface view. White represents hydrophilic residues whereas red represents hydrophobic residues. Zn ions indicated as silver spheres; water indicated as blue sphere.

We modeled the highest catalytic efficiency substrate for GloB, 1O, onto our structure. Autodock places 1O with its carbonyl oxygen adjacent to the active site water (Figure 5c)^42,43^. The GloB active site channel appears moderately wide, explaining why extensively branched substrates are not tolerated. Toward the end of the active site channel, GloB appears to form a tunnel. This tunnel is not reached by substrate 1O, but presumably would be occupied in more sterically bulky substrates such as 11O. One arm of this tunnel is comprised of the highly flexible loop, which is only partially visible in our electron density, suggesting that during catalysis this flexible loop may accommodate larger substrates, such as 11O.

### Esterase specificity of human and mouse sera

We sought to evaluate whether ester promoieties could be designed for microbe-specific activation. Using the same 32-compound fluorescent substrate library, we determined each substrate’s serum half-life. Both reconstituted and fresh sera functions are comparable in their activity and substrate preferences (Supplementary Figure 9).

We performed full kinetic profiling of lyophilized human sera to define its ester substrate specificity. As sera is a mixture of multiple proteins instead of a single protein species, we report the *V*_max_/*K*_m_ normalized to the total amount of protein added to the assay (Supplementary Figure 10A). As observed for FrmB and GloB (Supplementary Figures 10b, c), human sera has highest activity for oxygen and sulfur ether compounds. In contrast to FrmB and GloB, human sera is relatively uniform in its esterase activity across the substrate library. Short-chain substrates exhibit the highest catalytic efficiency, and though branching slightly reduces efficiency, it does not have as profound an impact as was observed with FrmB. The substrates displaying the poorest catalytic efficiency are universally the carbon series, in which added branching resulting in decreased substrate utilization.

As murine models are frequently used in the development and testing of novel pharmaceuticals, we also characterized the substrate preferences of mouse sera. Notably, mice are well known for their extremely active and broad serum esterase activity. Indeed, we find that mouse sera exhibit ~100-fold higher activity per mg serum protein than human sera (Supplementary Figure 10d, 11). This increase in activity is not uniform across the substrate library. Human sera is not as active on the carbon series and do comparatively better on the oxygen and sulfur ether compounds (Supplementary Figure 11b). Thus, our data suggest that mouse sera poorly predict human serum stability of ester prodrugs.

Finally, we sought to evaluate whether GloB and FrmB substrate specificities could be used for microbe-targeted prodrug design. As each esterase is likely to encounter multiple potential substrates *in vivo*, we utilized our modified *V*max/*K*m as a comparator. We performed pairwise analysis for each combination of FrmB and GloB against human and mouse sera (Figure 6a-d). Using a cutoff of 2^10^-fold enrichment in activity for the microbial enzymes over the serum enzymes, FrmB displays a preference over human sera for two compounds-3C and 6C, whereas GloB displays a preference for six compounds-2S, 3C, 10C, 11C, 11O, and 11S (Figure 6e). Conversely, mouse sera hydrolyze all compounds within this cutoff. Lowering the cutoff to a 2^5^-fold enrichment in catalytic efficiency over mouse sera, FrmB and GloB are both more specific for compound 2S, and GloB additionally displays specificity for compound 11O.

**Figure 6.**
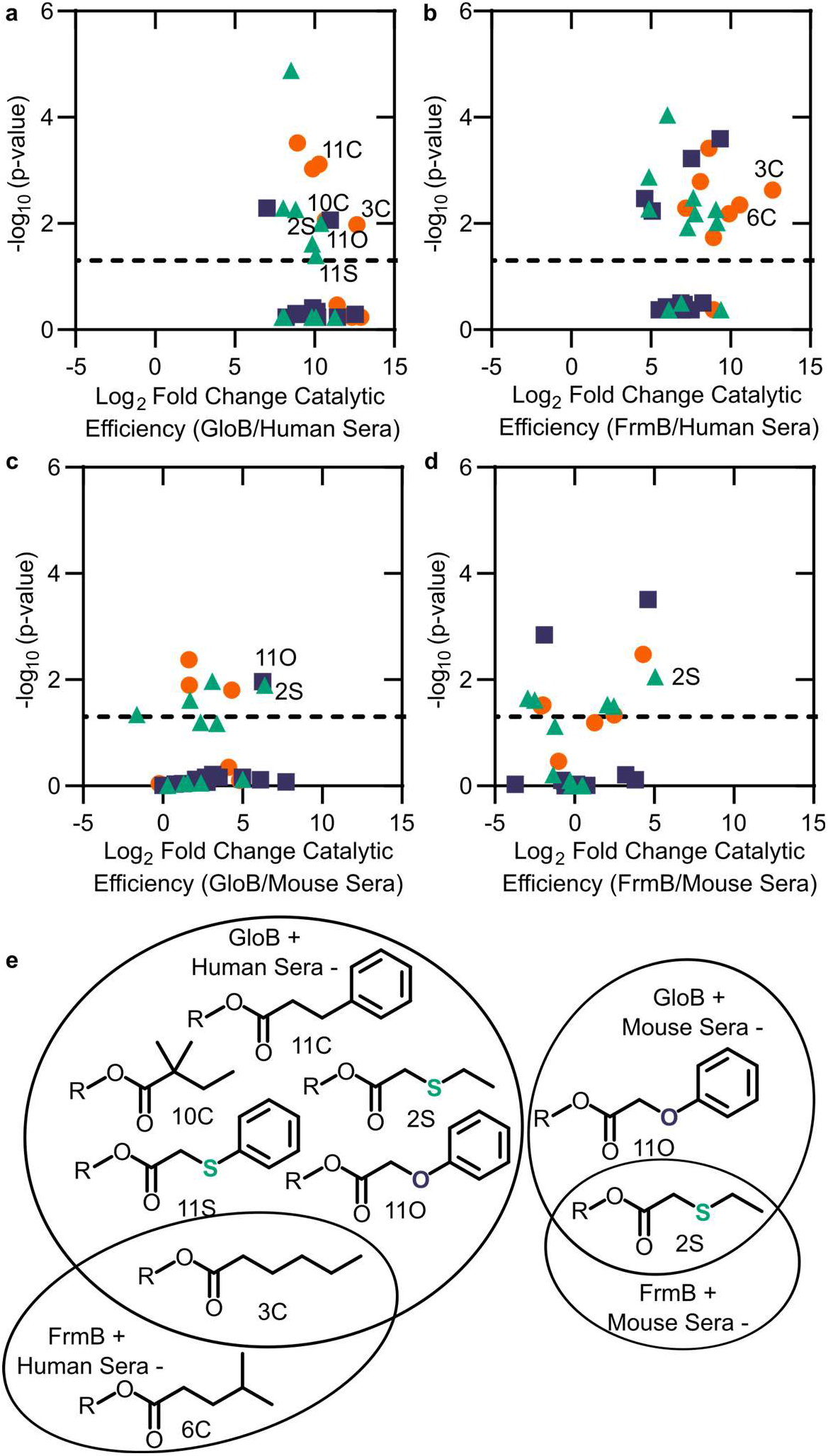
Comparison between microbial esterase and serum esterase catalytic efficiency. (a-d) Volcano plots of catalytic efficiency. Displayed are the means of three independent experiments. P-values calculated as pairwise t-tests with Holm-Sidak correction for multiple comparisons. (a) Comparison between human sera and GloB, (b) Human sera and FrmB, (c) Mouse sera and GloB, and (d) Mouse sera and FrmB). (e) Structures of ester substrates with 2^10^ enrichment in catalytic efficiency for microbial esterases over human serum (left), or 2^5^ enrichment over mouse serum. Dashed line indicates a p-value of 0.05.

## Discussion

Targeted microbial delivery and activation of lipophilic ester prodrugs is a highly desirable strategy to enable the expansion of druggable targets within bacteria while simultaneously improving drug selectivity. Identification of microbe-specific promoieties is crucial to this goal. Here, we identify two discrete esterases in *S. aureus*, FrmB and GloB, that activate the carboxy ester prodrug, POM-HEX (Supplementary Figure 12). Sequence homologues of FrmB and GloB are found broadly within microbes, though substantial sequence variation exists. FrmB and GloB have distinct ester substrate specificities, which are supported by the structures of their active sites. Importantly, enzymatic substrate specificity correlates with the rate of ester activation in live bacterial cells. Accordingly, simple modifications to ester prodrugs are sufficient to change their rates of activation *in vivo*.

Simple ester modifications can also change the pattern of prodrug activation. For the successful development of microbially targeted ester prodrugs, compounds must resist human enzymes. Here we demonstrate that human sera have specific ester substrate preferences, which are highly distinct from mouse sera, suggesting that rodents are an inadequate model for ester prodrug activation and therapeutic efficacy in humans. We find that the staphylococcal esterases FrmB and GloB utilize distinct patterns of substrates versus human sera. How microbes beyond *S. aureus* activate prodrugs, as well as the substrate specificities of pathogenic and commensal microbes, remains an important open question that will dictate the antimicrobial spectrum of a given ester prodrug. This work paves the way for structure-guided development of *S. aureus*-specific prodrugs and establishes a pipeline for the identification of additional microbial prodrug activating enzymes. We anticipate that these approaches will both guide the development of novel antimicrobials and lead to a more robust arsenal of anti-infective compounds with targeted specificity for the microbe over the human host.

## Methods

### Materials

POM-HEX, Hemi-HEX, and HEX were synthesized and resuspended in 100% DMSO as described previously^12^. Fluorescent ester compounds were generously provided by the laboratory of Geoffrey Hoops (Butler University), and synthesis and characterization has been previously described ^36^. Pooled, delipidated, defibrinated, and lyophilized human and mouse serum was obtained from Rockland Inc.

### Quantification of resistance

Half-maximal inhibitory concentration (IC_50_) determination was performed using microtiter broth dilution in clear 96-well plates^48^. Briefly, POM-HEX was added to 75 μL LB media at a final concentration of 20 – 50 μM POM-HEX and 0.5% DMSO, with POM-HEX concentrations varying according to resistant strain. Subsequently, POM-HEX was serially diluted in LB media containing 0.5% DMSO for a total of 10 dilutions. Two wells were left without drug, one used to define 100% growth, and the other used to control for media contamination and to define 0% growth. 75 μL of mid-log phase *S. aureus* diluted to 1 × 10^5^ colony forming units/mL were subsequently added to the plate. Following inoculation, plates were incubated at 37°C with shaking, and OD_600_ measurements were taken every 20 min for a total of 16 h. Half-maximal inhibitory concentrations (IC_50_) were determined by fitting the OD_600_ of each condition following 10 h of growth to a nonlinear regression using GraphPad Prism software. Experiments were performed in triplicate with technical duplicates.

### Generation of POM-HEX resistant strains

was performed by plating log-phase *S. aureus* Newman on LB agar containing 3.33 μM POM-HEX and incubating at 37°C overnight. Surviving single colonies were grown overnight in LB media and frozen in 10% glycerol for long term storage. All assays were performed from fresh overnight inoculations from glycerol stocks.

### Whole genome sequencing and variant discovery

Genomic DNA integrity was determined using Agilent 4200 Tapestation. Library preparation was performed with 0.25-0.5 μg of DNA. DNA was fragmented using a Covaris E220 sonicator using peak incident power 175, duty factor 10%, cycles per burst 200 for 240 sec at 4°C. DNA was blunt ended, had an A base added to the 3’ ends, and then had Illumina sequencing adapters ligated to the ends. Ligated fragments were then amplified for 9 cycles using primers incorporating unique dual index tags. Fragments were sequenced on an Illumina MiSeq using paired-end reads extending 150 bases.

For the analysis, the WT-assembled genomic sequence for *S. aureus* Newman was retrieved from GenBank (accession number AP009351.1), and the paired-end reads were aligned to this genome using Novoalign v3.03 (Novocraft Technologies). Duplicates were removed and variants were called using SAMtools^49^. SNPs were filtered against the sequenced parental strain, and genetic variants were annotated using SnpEff v4.3^50^. Whole genome sequencing data is available in the NCBI BioProject database and Sequence Read Archive under the BioProject ID PRJNA648156. All identified variants were verified via Sanger sequencing using the gene specific primers found in Supplementary Table 9. Amplicons were sequenced by GeneWiz Inc (Bejing, China).

### Recombinant expression and purification of FrmB and GloB

WT *frmB* and *gloB* sequences from *S. aureus* Newman were cloned into the BG1861 vector by GeneWiz Inc to introduce a hexahistidine tag^51^. The resultant plasmids were transformed into Stellar chemically competent cells (Clontech Laboratories), selected with carbenicillin, and the sequence was confirmed by Sanger sequencing. Subsequently, plasmids were transformed into chemically competent BL21 (DE3) cells and selected with 50 μg/mL ampicillin. Overnight liquid cultures were diluted 1:500 into LB media supplemented with ampicillin, grown shaking at 220 rpm to an OD_600_ of 0.5-0.8 at 37°C, chilled to 16°C and induced with 0.5 mM isopropyl β-D-1-thiogalactopyranoside (IPTG) for 16-20 h. Cells were harvested by centrifugation at 6,000 x g for 10 min at 4°C. The cell pellet was lysed by sonication in 50 mL lysis buffer containing 25 mM Tris HCl (pH 7.5), 250 mM NaCl, 20 mM imidazole, 1 mM MgCl_2_, 10% glycerol, and 200 μM phenylmethylsulfonyl fluoride (PMSF). Cell debris were removed by centrifugation twice at 20,000 x g for 20 min. The hexahistidine-tagged proteins were affinity purified from soluble lysate using nickel agarose beads (Gold Biotechnology). Bound protein was washed with 50 mL lysis buffer before elution using 5 mL of elution buffer containing 25 mM Tris HCl pH 7.5, 250 mM NaCl, 300 mM imidazole, 1 mM MgCl_2_, 10% glycerol. Affinity purified protein was further purified over a HiLoad 16/60 Superdex 200 gel filtration column (GE Healthsciences) using an AKTA Explorer. FPLC buffer contained 25 mM Tris HCl pH 7.5, 250 mM NaCl, 1 mM MgCl_2_, and 10% glycerol. Fractions containing >90% pure protein (evaluated by SDS-PAGE) were concentrated using an Amicon Ultra-15 centrifugal unit (EMD Millipore) and flash frozen in liquid nitrogen before storage at −80°C.

Protein used during crystallography experiments was generated via the same *frmB* and *gloB* sequences, but expression was performed from vector pET28a. *frmB* was cloned into the pET28a vector by GeneWiz Inc (Beijing, China) and *gloB* was cloned from the BG1861 vector using the forward primer 5’-dTGCTCGAGTGCGGCCGCTTAACCGTGTAAAAATGGATTT3’ and the reverse primer 5’-dCGCGCGGCAGCCATATGATGAGGATTTCAAGCTTAACTTT-3’. The PCR product was cloned into pET28a digested with restriction enzymes NotI and NdeI using InFusion HD Cloning (Takara Bio). Both cloning strategies introduce a hexahistidine tag followed by a thrombin cleavage sequence. The pET28a plasmids encoding either frmB or gloB were transformed into chemically-competent *E. coli* BL21(DE3) cells. Protein expression of FrmB proceeded as previously described, except FrmB containing cells were grown in Terrific Broth medium (24 g/L yeast extract, 20 g/L tryptone, 4 mL/L glycerol, 17 mM KH_2_PO_4_, 72 mM K_2_HPO_4_).

Selenomethionine (SeMet)-labeled GloB was prepared according to Van Duyne et al. with minor modifications^52^. Briefly, overnight cultures were grown in LB media, washed, and resuspended in M9 minimal media (per liter: 64 g Na_2_HPO_4_, 15 g KH_2_PO_4_, 2.5 g NaCl, and 5 g NH_4_Cl) supplemented with 50 mg EDTA, 8 mg FeCl_3_, 0.5 mg ZnCl_2_, 0.1 mg CuCl_2_, 0.1 mg CoCl_2_, 0.1 mg H_3_BO_3_, 16 mg MnCl_2_, 0.1 mg Ni_2_SO_4_, 0.1 mg molybdic acid, 0.5 mg riboflavin, 0.5 mg niacinamide, 0.5 mg pyridoxine monohydrate, and 0.5 mg thiamine per liter. Resuspended cultures were grown overnight. The following day, cultures were back diluted 1:50 and grown to an OD_600_ of 0.5-0.8 at 37°C. Once at the appropriate OD_600_ nm, the following amino acids were added to the culture media at: 100 mg/L: lysine, phenylalanine, and threonine, 50 mg/L: isoleucine, leucine, and valine, 60 mg/L: SeMet. Cultures were grown for an additional 15 min at 37°C before cells were chilled to 16°C and induced with 0.5 mM IPTG for 16-20 h.

Protein purification of FrmB and SeMet-labeled GloB for crystallography proceeded as previously except following affinity purification the elution was dialyzed for 16-20 h at 4°C with 20 U thrombin protease (GE Healthsciences) to remove the hexahistidine tag. Dialysis buffer contained 50 mM Tris pH 7.5, 50 mM NaCl, and 1 mM MgCl_2_. Following dialysis, uncleaved protein, the hexahistidine tag, and thrombin were removed by flowing dialyzed protein over a benzamidine sepharose and nickel agarose bead column (GE Healthsciences). Column flow through was further purified over a HiLoad 16/60 Superdex-200 gel filtration column (GE Healthsciences) equilibrated with dialysis buffer. Protein was concentrated to 8-10 mg/mL in an Amicon Ultra-15 centrifugal unit and frozen at −80°C.

### Fresh human serum

was collected from a willing volunteer in untreated BD vacutainer tubes (BD, BD366430) (Washington University IRB # 201012782). Whole blood was allowed to clot at room temperature, and aggregates were separated from the remaining serum through centrifugation at 400 x g for 8 min. Sera were obtained from the same volunteer on two separate occasions.

### Microfluidics measurements on*S. aureus*

Overnight cultures of *S. aureus* were grown in LB media, back diluted 1:500, and grown to early exponential phase (OD_600_ 0.1-0.15), then washed in 1X PBS and loaded on a bacterial CellASIC ONIX2 microfluidic plate. Prior to cell loading, the microfluidics plate lines were flushed with 1X PBS + 1% DMSO or 10 μM fluorescent pro-substrate in 1X PBS + 1% DMSO, and the plate was preincubated at 37°C. The microfluidics plate was loaded onto a Nikon Ti-E inverted microscope (Nikon Instruments, Inc) equipped with a 100x Plan N (N.A. = 1.45) Ph3 objective, SOLA SE Light Engine (Lumencor), heated control chamber (OKO Labs) and an ORCA-Flash4.0 sCMOS camera (Hammamatsu Photonics). The GFP filter set was purchased from Chroma Technology Corporation. Cells were loaded into the chamber until a single field of view contained 50-150 cells or cell clusters. Following cell loading, 1X PBS was slowly flown through the flow cell (t = 0), and cells were observed in both phase and fluorescence microscopy for 10 min before the flow media was rapidly switched over a period of two min to 1X PBS containing 1% DMSO and 10 μM fluorescent prosubstrate before slow flow for the remainder of the experiment. Images were acquired every two min for a total of 44 min, and all experiments were undertaken at 37°C. The phase contrast exposure time was kept constant at 200 ms and the fluorescent channel exposure time was kept constant at 500 ms. For fluorescent images, the gain remained constant across all experiments. Image capture and analysis was performed using Nikon Elements Advanced Research software. Individual cells or clusters of cells were auto detected in the fluorescent channel using the intrinsic background fluorescence of each cell. Manual curation followed autodetection to remove debris or cells that did not stay within the field of view throughout the experiment. Fluorescent intensity for each individual cell or cluster of cells was measured through the duration of the experiment and normalized to the area of the identified cell to yield the mean fluorescent intensity. Background cell autofluorescence was corrected by subtracting the average fluorescence across all identified objects from t = 0 through t = 10. Each experiment was performed in duplicate, with >50 individual cells or clusters analyzed in each experiment.

### Protein crystallography, phasing, and data refinement

Crystals of *S. aureus* FrmB were grown at 16°C using vapor diffusion in 20 μL hanging drops containing a 1:1 mixture of protein (6 mg/mL) and crystallization buffer (0.1 M Tricine pH 7.7, 15% PEG6K, 2.5 M NaCl, 0.125% *n*-dodecyl-β-D-glucoside). Prior to data collection, crystals were stabilized in cryoprotectant (mother liquor supplemented with 20% glycerol) before flash freezing in liquid nitrogen for data collection at 100 K. Crystals of Se-Met labeled *S. aureus* GloB were grown at 16°C using vapor diffusion in 2 μL hanging drops containing a 1:1 mixture of protein (8 mg/mL) and crystallization buffer (0.1 M imidazole pH 6.9, 0.2 M ammonium sulfate, 0.1 M calcium chloride, and 21% PEG 8k). SeMet-labeled GloB crystals were stabilized in well solution supplemented with 15% glycerol and flash frozen in liquid nitrogen. All diffraction images were collected at beamline 19-ID of the Argonne National Laboratory Advanced Photon Source at Argonne National Laboratory. HKL3000 was used to index, integrate, and scale the data sets^53^. To phase the initial dataset of FrmB, molecular replacement was performed in PHASER using the x-ray crystal structure of a low-temperature active alkaline esterase (PDB ID: 4RGY) as a search model^37,54^. SeMet-labeled GloB was phased using the x-ray crystal structure of TTHA1623 from *Thermus thermophilus* HB8 (PDB ID: 2ZWR)^44^. Buccaneer was used to build both initial models, and subsequent, iterative rounds of model building and refinement used COOT and PHENIX respectively^55–57^. Data collection and refinement statistics are summarized in Supplementary Table 6. Atomic coordinates and structure factors of *S.* aureus FrmB (PDB: 7L0A) and *S. aureus* GloB (PDB: 7L0B) are deposited in the RCSB Protein Data Bank.

### Substrate Docking

GloB and FrmB structures were prepared for substrate autodocking using AutoDock Tools 1.5.7^42^. Metals and water molecules were removed from the crystal structure of FrmB as canonical serine hydrolases do not utilize metal in their reaction mechanism. Solvent water in the GloB crystal structure was removed, but the active site water and heavily coordinated zinc molecules were left in place. The three-dimensional structure of substrate 1O was generated using ChemDraw3D, and prepared for docking using AutoDock Tools 1.5.7. Substrate docking of FrmB and GloB was performed using AutoDock Vina^43^.

## Supporting information

WT_S.aureus_3C_whole_cell_activation_movie

WT_S.aureus_5O_whole_cell_activation_movie

WT_S.aureus_1O_whole_cell_activation_movie

WT_S.aureus_9C_whole_cell_activation_movie

## Acknowledgements

We are grateful to the students of the Spring 2020 Biol 4522 course at Washington University for the creation of FrmB point mutation plasmids and the purification of FrmB point mutants. Thank you to Vandna Kukshal and Jason Schaffer for helpful discussions around data and mechanisms and to Petra Levin for assistance with microfluidics and microscopy. Fluorescent ester compounds were generously provided by the laboratory of Geoffrey Hoops (Butler University). A.O.J is supported by NIH/NIAID R01-AI103280, R21-AI123808, and R21-AI130584, and AOJ is an Investigator in the Pathogenesis of Infectious Diseases (PATH) of the Burroughs Wellcome Fund. This publication was made possible in part by Grant Number UL1 RR024992 from the NIH-National Center for Research Resources (NCRR)

