## Supplemental Methods, Figures, and Tables for "Structure-guided microbial targeting of antistaphylococcal prodrugs"

[1]Department of Pediatrics, Washington University School of Medicine, St. Louis, MO, 63110, USA; [2]Department of Biology, Washington University in St. Louis, St. Louis, MO, 63130, USA; [3]Department of Cancer Systems Imaging, The University of Texas MD Anderson Cancer Center, Houston, TX, 77054, USA; [4]Department of Pediatrics, Perelman School of Medicine, University of Pennsylvania, Children's Hospital of Philadelphia, Philadelphia, PA, 19104, USA; [5] Department of Chemistry, The George Washington University, Washington DC, 20052, USA [6]Department of Molecular Microbiology, Washington University School of Medicine, St. Louis, MO, USA;

**WhatsGNU analysis.** The *S. aureus* database was used to produce WhatsGNU proteomic reports for all the strains using WhatsGNU_main.py script in the ortholog mode. Eighteen *S. aureus* (9 atopic dermatitis (AD) and 9 soft and skin tissue infection (SSTI)) isolates from an ongoing project representing different clonal complexes (CC1/5/8/22/30) were used for the comparison. The CC details for the 18 isolates are provided in Supplementary Table 10. The reports were then used to produce a heat map of the GNU scores of GloB and FrmB using the heat map function in the WhatsGNU_plotter.py script. The heatmap was annotated with the ortholog variant rarity index where 'r' represents a rare GNU score (in the context of other alleles in the same protein ortholog group).

**Phylogenetic tree construction.** Sequences of GloB, FrmB, and RpoB orthologs were retrieved from NCBI using the BlastP function with each organism on the tree as an individual search set. Of the returned sequences, the first complete sequence with the lowest E-value was selected for further analysis. Organisms were selected to include a wide variety of pathogenic and commensal microbes[1]. In one instance, several of the top *E. coli* sequences were found to be highly similar to *S. aureus*, and on further analysis we discovered that the original sequencing samples had high levels of *S. aureus* reads. These contaminated sequences were disregarded in our analysis. Sequence alignment was performed using MUSCLE using the default parameters, and the unrooted phylogenetic trees were visualized using iTOL[2,3].

**Glyoxalase II activity assay.** Glyoxalase II activity was assessed as previously with minor changes[4,5]. Briefly, reactions were mixed to form a final concentration of 25 mM Tris pH 7.5, 250 mM NaCl, 1 mM $MnCl_2$, 10% glycerol, 200 μM 5,5'-dithiobis(2-nitrobenzoic acid) (DTNB, Sigma D8130), 1 mM D-lactoylglutathione (Sigma L7140) and 0.15-0.63 μg protein (130-550 nM GloB, 100-430 nM FrmB). Protein concentrations were varied to ensure the reaction was linear across protein concentrations. Reactions without D-lactoylglutathione were pre-incubated at 37ºC for 10 min prior to assay initiation with the addition of substrate. Release of glutathione from D-lactoylglutathione was quantified spectrophotometrically at 37ºC and 412 nm through the conversion of DTNB to TNB. Experiments were performed in triplicate with technical duplicates.

**4-nitrophenyl ester substrate activity assays.** 4-Nitrophenyl substrate specific activity was determined in 50 µL reactions containing 25 mM Tris pH 7.5, 250 mM NaCl, 1 mM $MnCl_2$, 10% glycerol, 1 µM protein, and 1 mM 4-nitrophenyl substrate. The tested substrates, 4-nitrophenyl acetate (Sigma, N8130), 4-nitrophenyl butyrate (Sigma N9876), and 4-nitrophenyl trimethylacetate (Sigma 135046) were resuspended in acetonitrile at 100 mM. Reactions without 4-nitrophenyl substrate were preincubated at 37ºC for 10 min prior to assay initiation via substrate addition. Conversion of 4-nitrophenyl substrates to 4-nitrophenol was tracked photometrically at 37ºC and $A_{405nm}$. Experiments were performed in triplicate with technical duplicates.

**NMR characterization of GloB and FrmB activation products.** 200 or 400 µM POM-HEX was incubated with 4 nmol GloB, FrmB, or 4 nmol each GloB and FrmB in 500 µL reactions. Reactions were buffered to a final concentration of 50 mM Tris pH 7.5, 50 mM NaCl, 1 mM $MgCl_2$. Reactions were allowed to proceed for 1 h at 37˚C prior to analysis. Samples were prepared for NMR studies by resuspending them in water and 10% (50 µL) $D_2O$ (deuterium oxide 99.9% D, contains 0.75 wt% 3-(trimethylsilyl)propionic-2,2,3,3-$d_4$ acid, sodium salt, Sigma–Aldrich). NMR spectra are acquired on a Bruker Avance III HD 500 MHz spectrometer equipped with a cryoprobe. Two-dimensional (2D) $^{1}H$-$^{31}P$ heteronuclear single quantum correlation (HSQC) measurements were obtained using hsqcetgp pulse program (with duration of 15 min and scan parameters of 2 scans, td=1024 and 256, gpz2 %=32.40, $^{31}P$ SW= 40 ppm, O2p=20 ppm, cnst2=22.95) and analyzed using 3.1 TopSpin. The 1D projection of columns excluding the water signal was obtained from the 2D $^{1}H$-$^{31}P$ HSQC spectrum by obtaining spectra of positive projection of columns 1 to 600 and 650 to 1024 and adding them.

**Esterase substrate specificity determination using fluorogenic SAR library.** Kinetic measurements were performed according to White *et al.* with minor variation[6]. Lyophilized human and mouse sera were resuspended according to manufacturer instructions in highly pure, filtered water at protein concentrations of 85 mg/mL and 70 mg/mL, respectively. 1 mL of resuspended serum was added to a 24 mL mastermix for a final concentration of 31.25 mM Tris pH 7.5, 312.5 mM NaCl, 1.25 mM $MgCl_2$, 12.5% glycerol, and 3.4 mg/mL or 2.8 mg/mL protein for human and mouse serum, respectively. For purified proteins, 5 mL of a 75 µg/mL stock was added to yield a 20 mL mastermix containing 31.25 mM Tris pH 7.5, 312.5 mM NaCl, 1.25 mM $MgCl_2$, 12.5% glycerol, and 18.75 µg/mL protein. Mastermix was stored on ice when not in use. 20 µL of mastermix was transferred to a black, 96-well half area microplate (Corning, CLS3993) and prewarmed at 37°C. Fluorogenic substrates were prepared as 10 mM stock solutions in 100% DMSO and were diluted in water to a starting concentration of 500 µM. Enzyme catalyzed substrate hydrolysis was initiated by addition of 5 µL substrate dilution in technical duplicate to the prewarmed serum or protein solution. Final assay concentrations were: 25 mM Tris pH 7.5, 250 mM NaCl, 1 mM $MgCl_2$, 10% glycerol, and protein at a concentration of 2.72 mg/mL (human serum), 2.24 mg/mL (mouse serum), or 15 µg/mL (FrmB, GloB). The resulting change in fluorescence ($\lambda_{ex}$ = 485 nm, $\lambda_{em}$ = 520 nm) was followed for 15 min at 37°C, collecting data every 30 sec on a FLUOstar Omega microplate reader (BMG Labtech). Fluorescence measurements were converted to molar concentrations using a fluorescein standard curve (2.5 nmol-0.6 pmol). The initial rates of reaction were measured three independent times with two technical replicates per measurement and fit to a line using GraphPad Prism (GraphPad Software, La Jolla, CA). Initial rates of reaction were plotted versus the concentration of substrate and fit to a standard Michaelis-Menten equation ($v = V_{max}[S]/(K_m + [S])$), yielding estimates of $V_{max}$ and $K_m$. Values for $k_{cat}$ and $k_{cat}/K_m$ were calculated based on amount of enzyme added when purified enzymes were used. For substrates that did not display saturation in the velocity versus substrate experiment (i.e., $K_m$>>>[S]), $V_{max}/K_m$ was estimated based on v/[S].

**Serum half-life determination.** Lyophilized human sera was obtained from Rockland Inc. and resuspended in pure water. 20 µL lyophilized sera or fresh sera was prewarmed at 37ºC in a 96-well half-area microplate (Corning, CLS3993). Following plate warming, 5 µL of the fluorogenic substrates were added to the plate for a final concentration of 25 µM. Substrate hydrolysis was tracked over a period of three h at 37ºC, with fluorescence measurements ($\lambda_{ex}$ = 485 nm, $\lambda_{em}$ = 520 nm) being taken every two min on a FLUOstar Omega microplate reader (BMG Labtech). The resulting fluorescence values were converted to % substrate hydrolyzed using a fluorescein standard curve and fit to a one-phase decay model using GraphPad Prism. Experiments were performed in technical and biological duplicate.

### Supplementary Figures

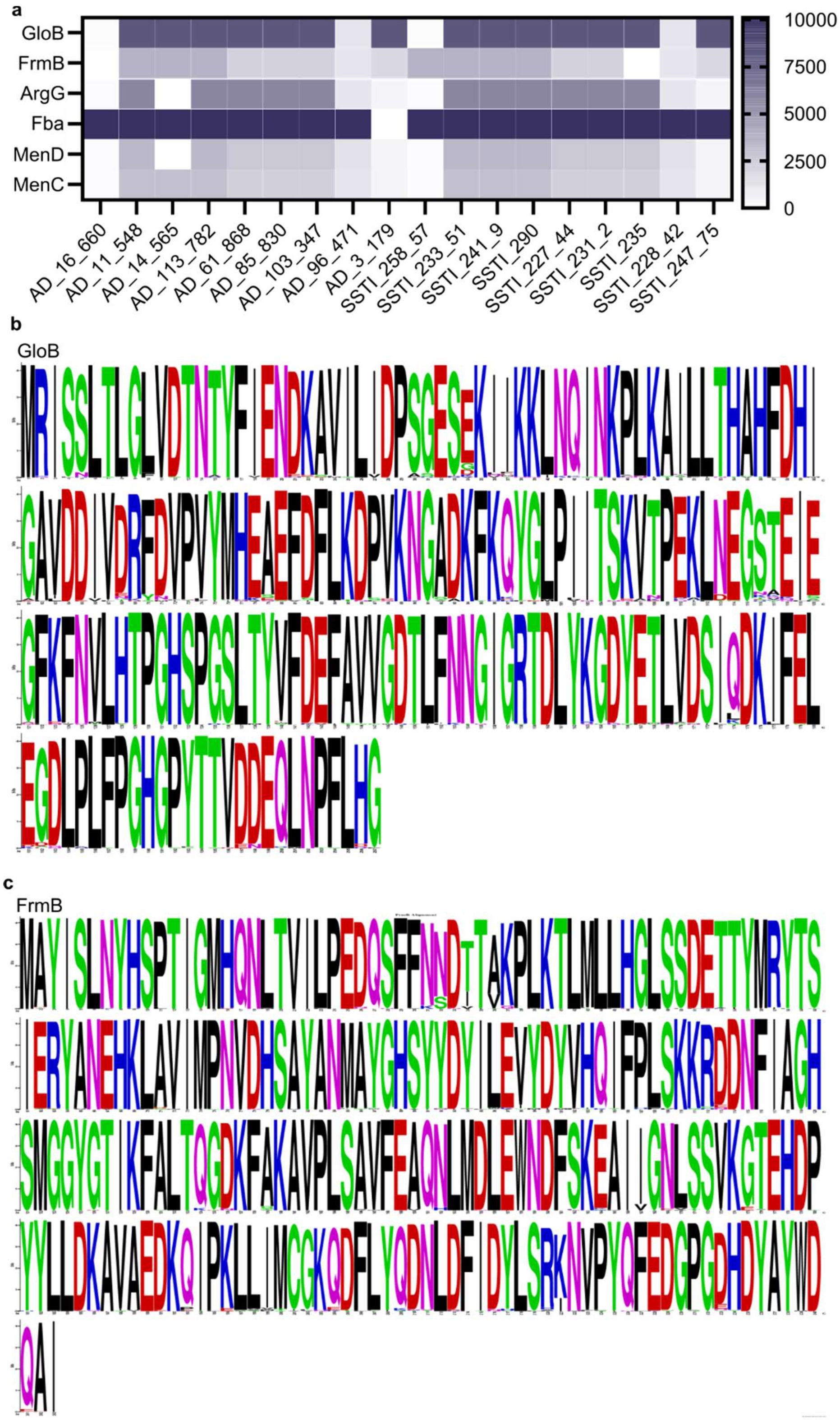

**Supplementary Figure 1** Conservation of FrmB and GloB within *S. aureus*. (a) WhatsGNU analysis of GloB and FrmB. Control proteins: ArgG, argininosuccinate synthase;, Fba, fructose-bisphosphate aldolase; MenD, 2-succinyl-5-enolpyruvyl-6-hydroxy-3-cyclohexene-1carboxylate synthase; and MenC, o-succinylbenzoate synthase. GNU stands for gene novelty unit and is a count of how many protein sequences in the database have an exact match to the queried sequence, with higher counts indicating sequence conservation. Strains across the x-axis are representative strains from the 18 *S. aureus* colony complexes which were used to query the *S. aureus* database. (b, c) MAFFT alignment of GloB (b) and FrmB (c) protein sequences across the *S. aureus* sequence database.

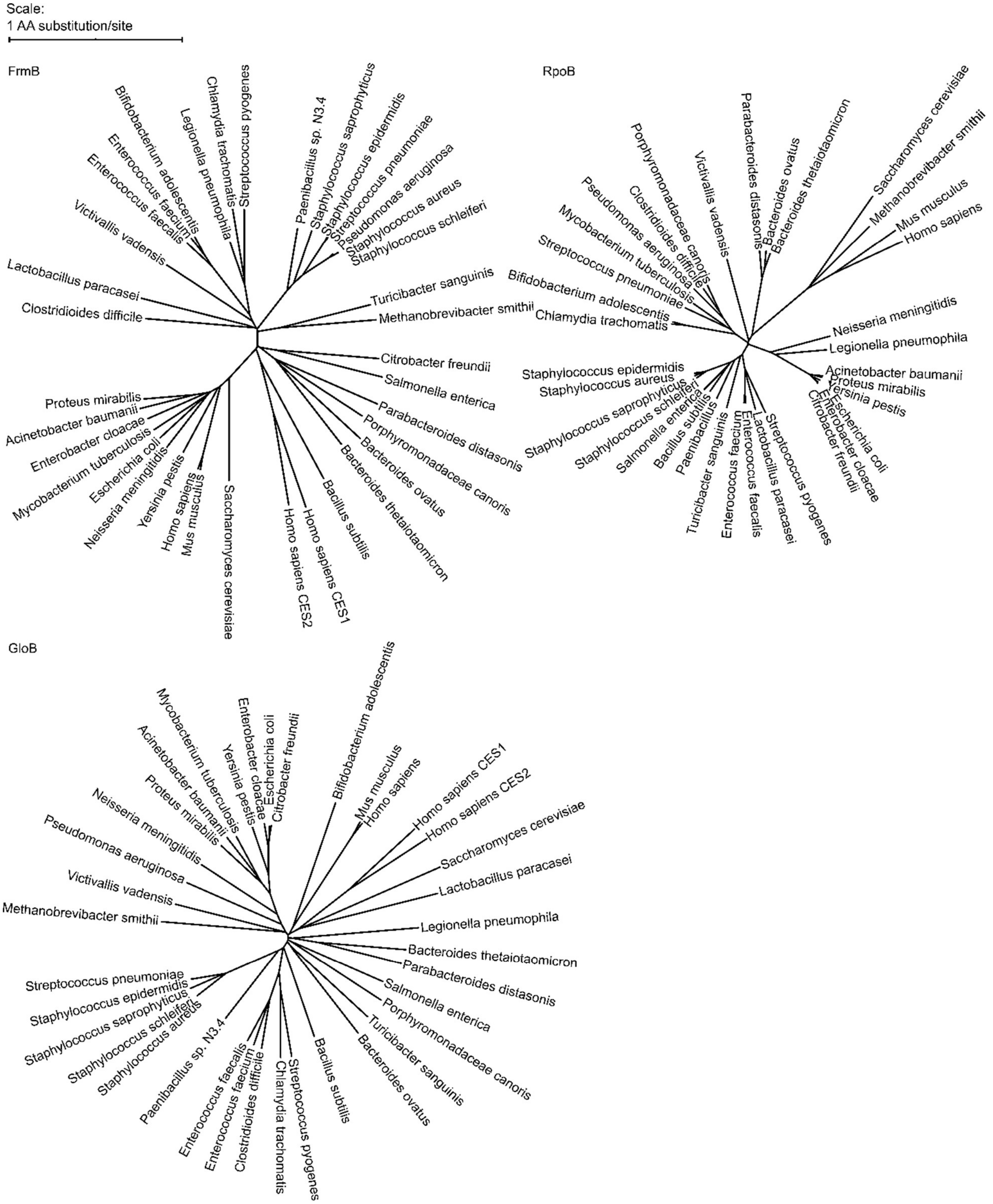

**Supplementary Figure 2** Phylogenetic tree of FrmB and GloB. Sequences of GloB, FrmB, and RpoB were retrieved from NCBI using BlastP against each organism. Sequence alignment performed using MUSCLE and alignment visualized using the interactive Tree of Life (iTOL).

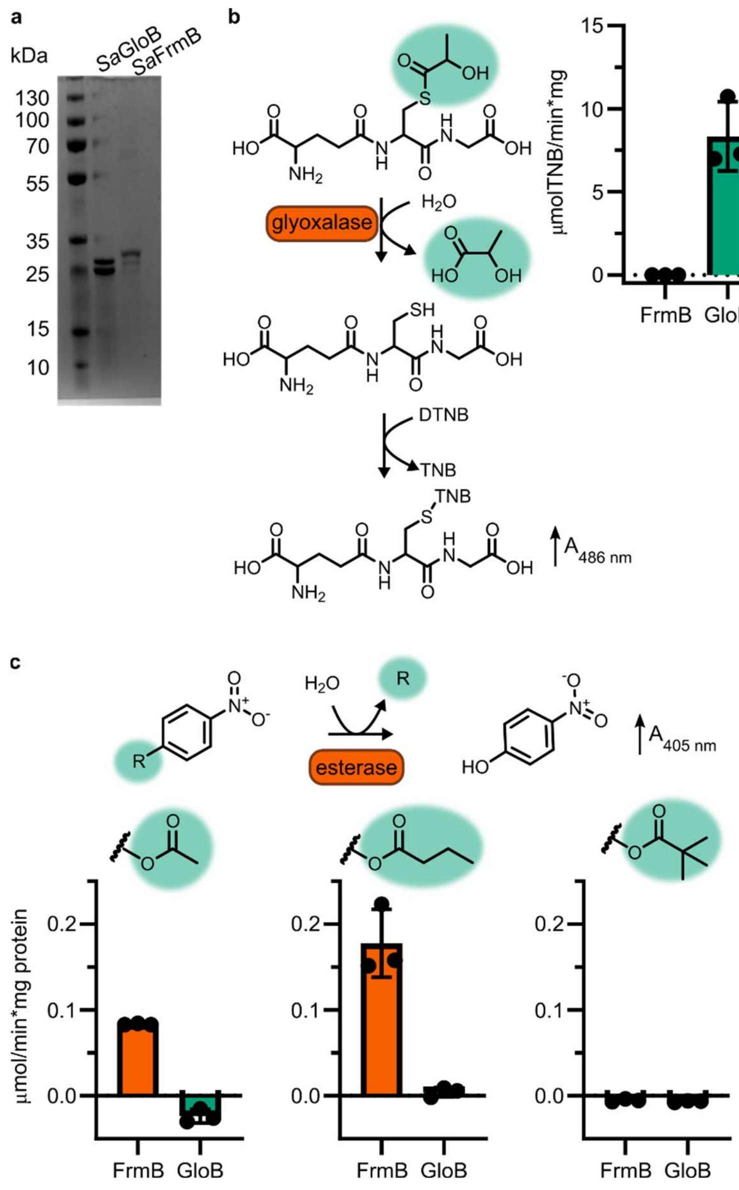

**Supplementary Figure 3** Enzymatic characterization of GloB and FrmB. (a) SDS-PAGE gel of GloB and FrmB protein preparations. Expected molecular weights are 23.3 kDa and 29.5 kDa, respectively. (b) Glyoxalase II activity assay, enzymatic conversion of S-lactoylglutathione releases free glutathione and reacts with DTNB resulting in increased absorbance at 412 nm. (c) 4-Nitrophenyl activation results in increased absorbance at 405 nm. Left to right, activity when supplied 4-nitrophenyl acetate, 4-nitrophenyl butyrate, and 4-nitrophenyl trimethyl acetate. Displayed in points is the mean of two technical replicates for individual experiments, bars indicate mean ± SD of three independent biological experiments performed in technical duplicate.

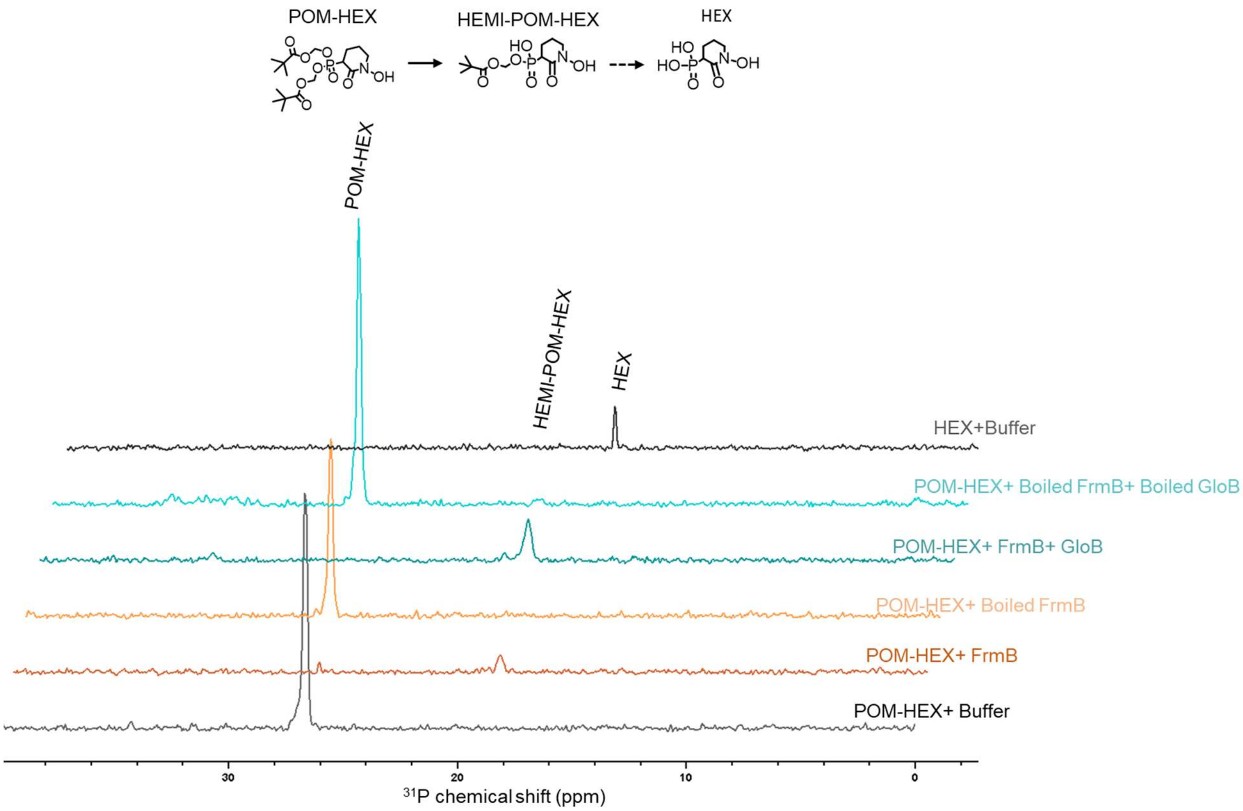

**Supplementary Figure 4** NMR characterization of POM-HEX activation by GloB and FrmB. Two-dimensional (2D) $^{1}H$-$^{31}P$ HSQC NMR spectra of products following incubation of FrmB, GloB, catalytically inactive (boiled) GloB and FrmB, or buffer alone. Also included are the $^{1}H$-$^{31}P$ HSQC NMR spectra of POM-HEX and HEX. Displayed are representative traces of three independent experiments. HEMI-POM HEX peak inferred by predicted shift.

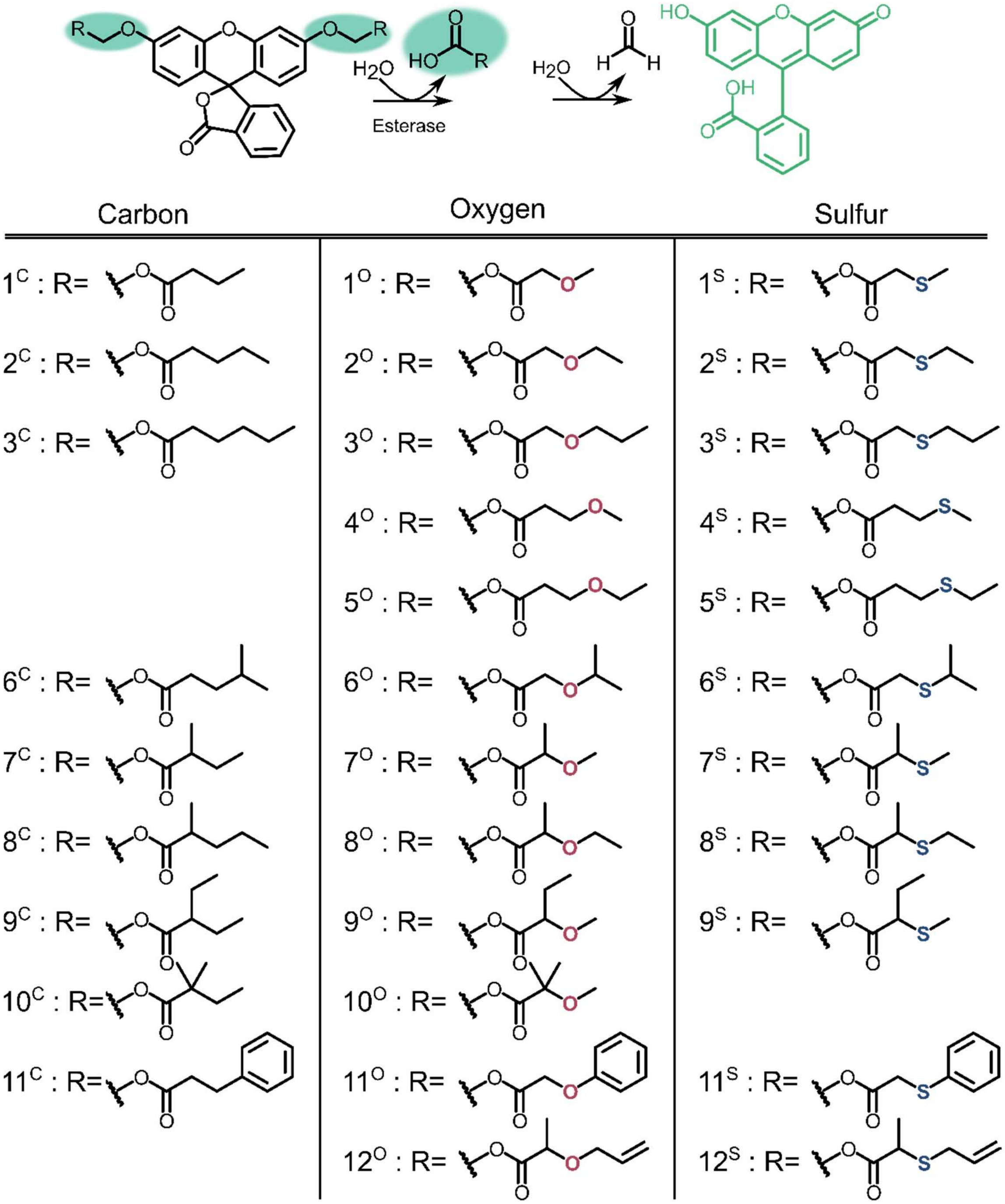

**Supplementary Figure 5** Profluorescent substrate library. Activation of substrates via esterase action results in fluorescence.

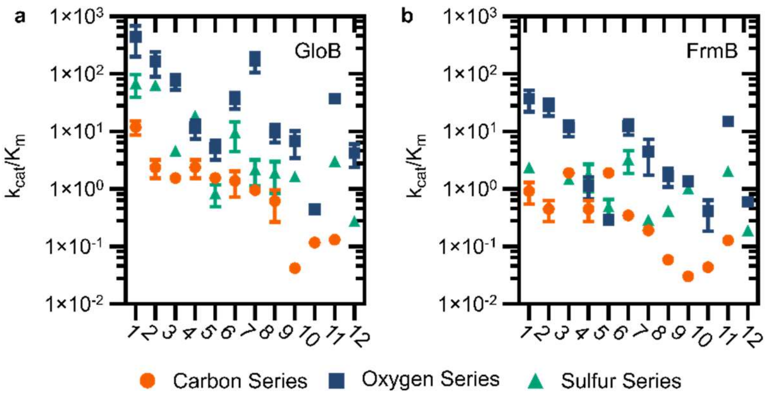

**Supplementary Figure 6** Catalytic efficiency of GloB and FrmB. Numbers correspond to the structures displayed in Figure S5, compounds in the carbon series denoted in orange, oxygen series in blue, and sulfur series in green.

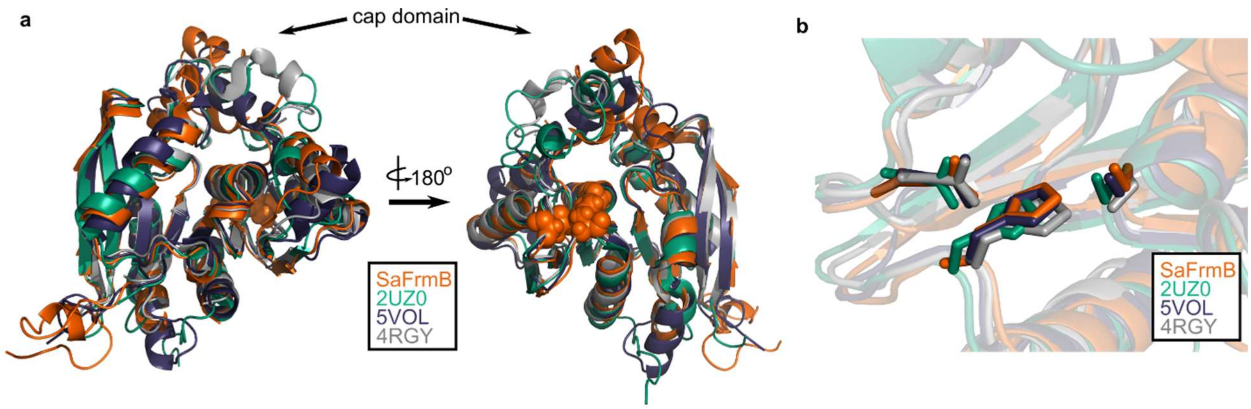

**Supplementary Figure 7** Structural conservation of FrmB. (a) Overall structural alignment of FrmB (orange) with *S. pneumonia* EstA (PDB:2UZ0), *B. intestinalis* ferulic acid esterase (PDB:5VOL), and deep sea bacteria Est12 (PDB4RGY). (b) Conservation of the serine hydrolase catalytic triad in FrmB and related proteins.

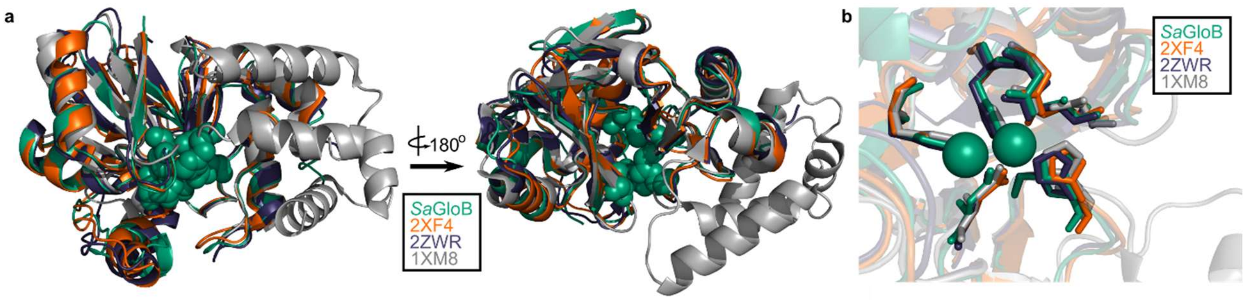

**Supplementary Figure 8** Structural conservation of GloB. (a) Overall structural alignment of GloB (green) with *S. enterica* YcbI (PDB:2XF4), *T. thermophilus* TTHA1623 (PDB:2ZWR), and *A. thaliana* glyoxalase II (PDB:1XM8). Zinc coordinating residues are colored in green spheres. (b) Positioning of the zinc coordinating residues (green spheres).

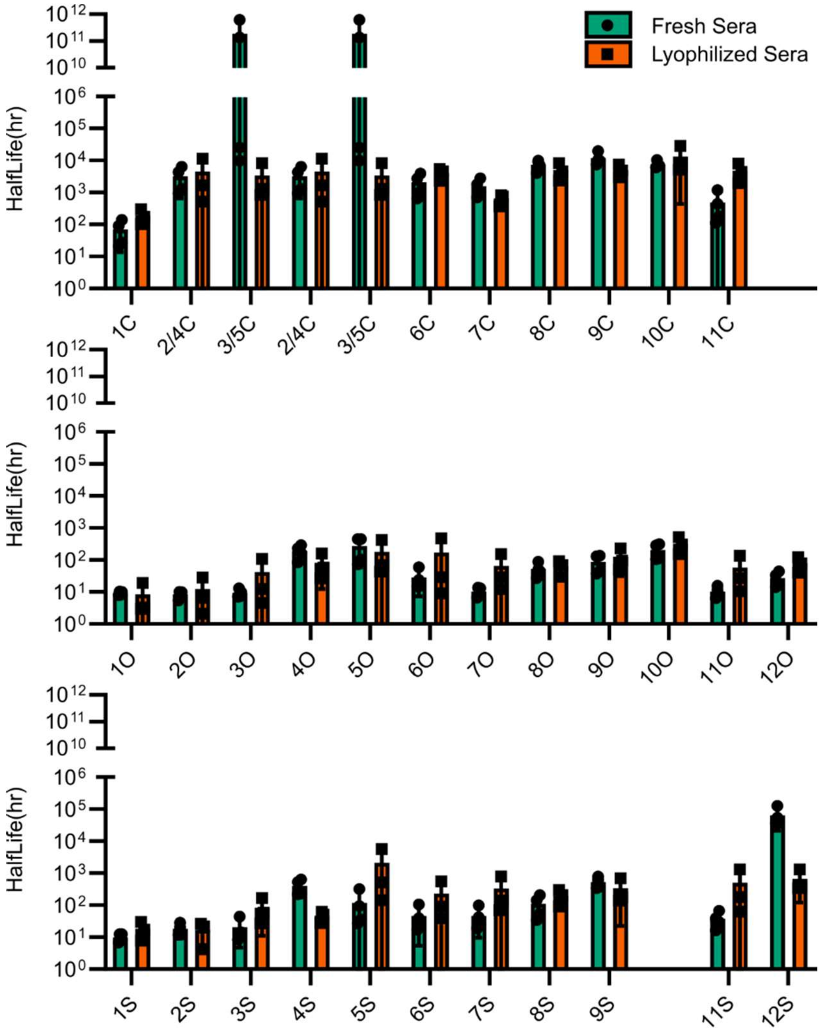

**Supplementary Figure 9** Comparison of esterase activity between fresh and lyophilized human sera. Points represent individual experiments; bars represent the mean ± SD of the four replicates.

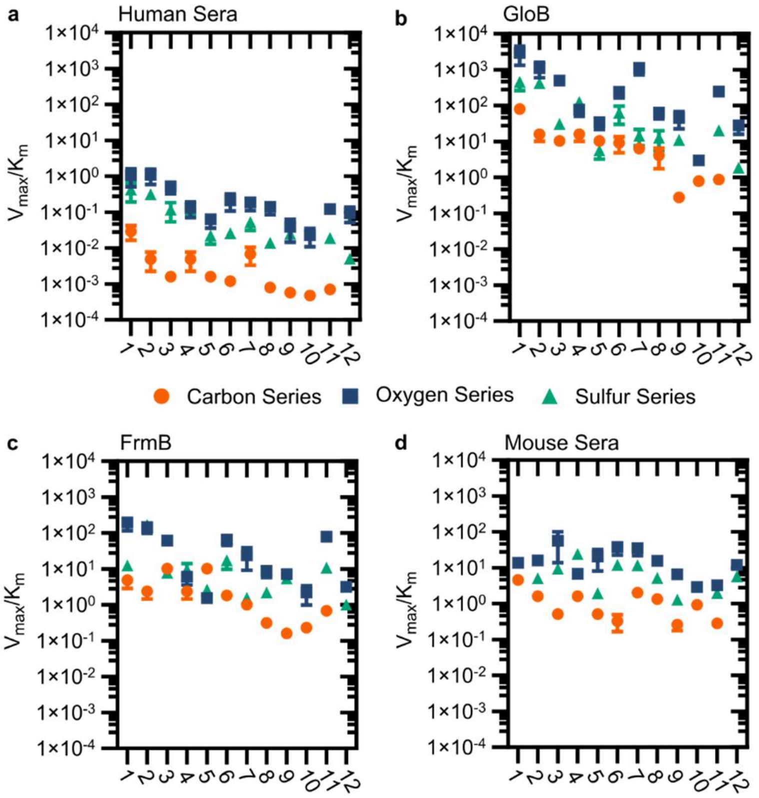

**Supplementary Figure 10** Modified vatalytic efficiency (pmol fluorescein produced * $min^{-1}$*$\mu g^{-1}$ protein) of human sera, GloB, FrmB, and mouse sera. X-axis corresponds to compound identities in Figure S5. Carbon containing compounds indicated in orange, oxygen in blue, and sulfur in green. Displayed are the means ± SD of three independent biological experiments.

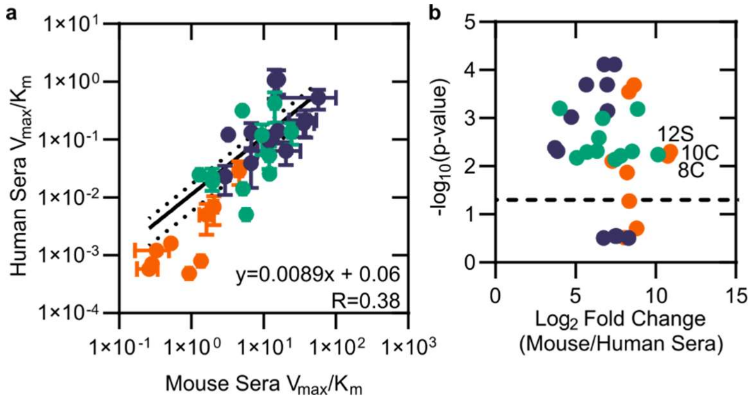

**Supplementary Figure 11** Comparison of mouse and human sera. (a) Modified catalytic efficiency (pmol fluorescein produced * $min^{-1}$*$\mu g^{-1}$ protein) of human and mouse sera. Displayed is a linear regression of the fit between mouse and human sera. (b) Volcano plot of catalytic efficiency. Displayed are the means of three independent experiments. P-values calculated as pairwise t-tests with Holm-Sidak correction for multiple comparisons. Dashed line indicates a p-value of 0.05.

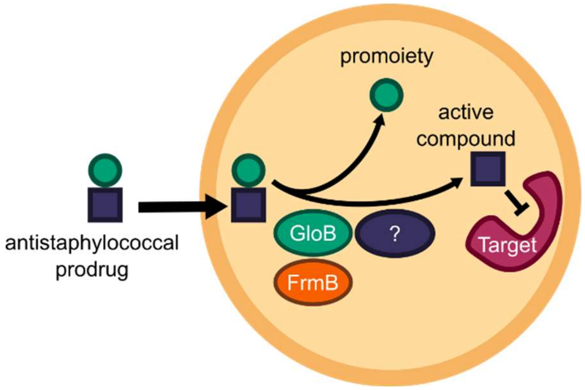

**Supplementary Figure 12** Model of antistaphylocccal prodrug activation. Lipophilic carboxy ester prodrugs transit the cell membrane, are first activated by either GloB or FrmB and at least one additional enzyme, before inhibiting the cellular target.

**Supplementary Table 1** Half-maximal inhibitory concentration ($IC_{50}$) values for POM-HEX against predicted prodrug activating esterases. $IC_{50}$ values are the result of three independent biological experiments with technical duplicates. P-values calculated as a one-way ANOVA with Dunnett's correction for multiple comparisons.

| SAUSA300 gene | Newman gene | StrID (NARSA) | Pan gene symbol | POM-HEX $IC_{50}$ (µM) | POM-HEX $IC_{50}$ SD (µM) | Significantly different from WT (JE2)? | Adjusted P-value | Predicted function | GO term |
|---|---|---|---|---|---|---|---|---|---|
| Parental Strain | | JE2 | | 1.54 | 0.40 | | | N/A | Parental Strain |
| SAUSA300_0742 | NWMN_0727 | NE145 | *uvrA* | 1.46 | 0.17 | ns | 0.9999 | Exonuclease ABC, A subunit | carboxylic ester hydrolase activity, carboxylesterase activity |
| SAUSA300_1902 | NWMN_1859 | NE202 | | 0.700 | 0.066 | ns | 0.9101 | Hypothetical protein | phosphoric diester hydrolase activity |
| SAUSA300_2173 | NWMN_2121 | NE223 | *truA* | 1.39 | 0.29 | ns | 0.9997 | tRNA pseudouridine synthase A | phosphatase |
| SAUSA300_1285 | NWMN_1303 | NE293 | | 1.41 | 0.64 | ns | 0.9997 | ABC transporter, ATP-binding protein | phosphatase |
| SAUSA300_2515 | NWMN_2477 | NE355 | *gbaA* | 1.73 | 0.48 | ns | 0.9997 | Transcriptional regulator, TetR family | carboxylic ester hydrolase activity |
| SAUSA300_1505 | NWMN_1449 | NE377 | *gloB* | 10.6 | 3.2 | **** | <0.0001 | Hydroxyacylglutathione hydrolase | hydroxyacylglutathione hydrolase |
| SAUSA300_0142 | NWMN_0084 | NE478 | *phnE* | 1.61 | 0.81 | ns | 0.9999 | Phosphonate ABC transporter, permease protein | phosphoric diester hydrolase activity |
| SAUSA300_2564 | NWMN_2528 | NE503 | *estA, frmB* | 6.28 | 2.25 | **** | <0.0001 | Tributyrin esterase | carboxylic ester hydrolase activity |
| SAUSA300_2473 | NWMN_2434 | NE532 | | 1.56 | 0.51 | ns | >0.9999 | Hypothetical Alkaline Phosphatase | phosphatase |
| SAUSA300_0581 | NWMN_0561 | NE621 | | 1.62 | 0.55 | ns | 0.9999 | Hypothetical protein | phosphatase activity |
| SAUSA300_0299 | | NE812 | | 1.64 | 0.56 | ns | 0.9998 | Hypothetical protein | phosphatase activity |
| SAUSA300_1993 | NWMN_1947 | NE937 | *fruC* | 1.67 | 0.52 | ns | 0.9998 | PfkB family kinase | phosphatase |
| SAUSA300_1752 | NWMN_1700 | NE949 | *hsdM2* | 1.73 | 0.53 | ns | 0.9996 | type I restriction-modification system, M subunit | phosphoric diester hydrolase activity |
| SAUSA300_0312 | NWMN_0254 | NE1039 | *psuG* | 1.73 | 0.51 | ns | 0.9997 | Hypothetical protein | carboxylic ester hydrolase activity |
| SAUSA300_0538 | NWMN_0515 | NE1071 | *capD* | 1.54 | 0.41 | ns | >0.9999 | NAD-dependent epimerase/dehydratase family protein | phosphatase activity |
| SAUSA300_1792 | NWMN_1735 | NE1173 | | 1.84 | 0.40 | ns | 0.9994 | Hypothetical protein | phosphatase activity |

| SAUSA300 gene | Newman gene | StrID (NARSA) | Pan gene symbol | POM-HEX $IC_{50}$ (µM) | POM-HEX $IC_{50}$ SD (µM) | Significantly different from WT (JE2)? | Adjusted P-value | Predicted function | GO term |
|---|---|---|---|---|---|---|---|---|---|
| SAUSA300_0421 | NWMN_0414 | NE1225 | | 2.15 | 0.48 | ns | 0.9938 | Hypothetical protein | carboxylic ester hydrolase activity, carboxylesterase |
| SAUSA300_0214 | NWMN_0156 | NE1238 | | 2.08 | 0.54 | ns | 0.9986 | Hypothetical protein | phosphoric diester hydrolase activity |
| SAUSA300_0690 | NWMN_0674 | NE1296 | *saeS* | 2.07 | 0.53 | ns | 0.9986 | Sensor histidine kinase SaeS | phosphatase |
| SAUSA300_1639 | NWMN_1586 | NE1486 | *phoP* | 2.18 | 0.41 | ns | 0.9932 | Alkaline phosphatase synthesis transcriptional regulatory protein | phosphoric diester hydrolase activity |
| SAUSA300_0840 | NWMN_0808 | NE1505 | | 2.20 | 0.54 | ns | 0.9927 | Hypothetical protein | phosphoric diester hydrolase activity |
| SAUSA300_1563 | NWMN_1507 | NE1519 | *accC* | 1.75 | 0.70 | ns | 0.9996 | Acetyl-CoA carboxylase, biotin carboxylase | phosphoric diester hydrolase activity |
| SAUSA300_2508 | | NE1547 | | 1.48 | 0.23 | ns | >0.9999 | Hypothetical protein | phosphatase activity |
| SAUSA300_0996 | NWMN_962 | NE1610 | *pdhD* | 1.23 | 0.81 | ns | 0.9993 | Dihydrolipoamide dehydrogenase | phosphoric diester hydrolase activity |
| SAUSA300_2367 | NWMN_2320 | NE1682 | *hlgB* | 1.37 | 0.19 | ns | 0.9997 | Gamma-hemolysin component B | phosphatase |

**Supplementary Table 2** Genotype and phenotype of POM-HEX resistant *S. aureus*. Displayed are the whole genome sequencing mutations that have been verified. Called mutations that were not observed via confirmatory Sanger sequencing are excluded. $IC_{50}$ values are the result of three independent biologic replicates with technical duplicates.

| **Gene (Newman)** | **WT_SAUR** | **R1** | **R2** | **R3** | **R4** | **R5** | **R6** | **R7** | **R8** | **R9** |
|---|---|---|---|---|---|---|---|---|---|---|
| NWMN_1449 (*gloB*) | | | | | | | c.376G>T<br>p.Val126Leu | | c.430T>A<br>p.Phe144Ile | c.433G>A<br>p.Ala145Thr |
| NWMN_2528 (*estA, frmB*) | | | | c.40G>C<br>p.Gly14Arg | | | | | | |
| NWMN_0144 | | | | | c.1472G>T<br>p.Arg491Met | | | | | |
| NWMN_0240 | | | | | | | | | | |
| NWMN_0309 | | | | c.23A>C<br>p.Asn8Thr | | c.23A>C<br>p.Asn8Thr | | | | c.23A>C<br>p.Asn8Thr |
| NWMN_0471 (*tilS*) | | | | | | | | | | |
| NWMN_0523 (*sdrC*) | | | | | | | | | | |
| NWMN_0564 | | | | | | | | | | |
| NWMN_0954 | | | | | | | | | | |
| NWMN_1152 | | | | | | | | | | |
| NWMN_1311 (*lysA*) | | | | | | | | | | |
| NWMN_1425 (*recN*) | | | | | | | | | | |
| NWMN_1655 | | | c.17G>A<br>p.Arg6His | c.17G>A<br>p.Arg6His | | | | | | |
| NWMN_1735 | | | | | | | | | | |
| NWMN_1754 | | | | | | | | | | |
| NWMN_2040 (*pdp*) | | | | | | | | | | |
| NWMN_2253 | | c.1412C>T<br>p.Ala471Val | c.1412C>T<br>p.Ala471Val | | | c.1412C>T<br>p.Ala471Val | | | | |
| NWMN_2388 | | | | | | | | | | |
| POM-HEX $IC_{50}$ (μM) | 1.921 | 6.337 | 5.932 | 9.561 | 10.54 | 3.752 | 11.09 | 10.71 | 16.96 | 17.71 |
| Std. Deviation | 1.111 | 0.8778 | 0.7325 | 0.4846 | 6.261 | 2.594 | 4.974 | 6.602 | 7.264 | 10.65 |

| Gene (Newman) | R10 | R11 | R12 | R13 | R14 | R15 | R16 | R17 | R18 |
|---|---|---|---|---|---|---|---|---|---|
| NWMN_1449 (*gloB*) | c.70G>T p.Val24Phe | c.166C>T p.His56Tyr | c.326C>A p.Pro109His | c.70G>T p.Val24Phe | c.70G>T p.Val24Phe | c.289C>T p.Gln97* | | | |
| NWMN_2528 (*estA*, *frmB*) | | | | | | | c.218_219insCATATGCCATGTTAGCA p.Met74fs | c.366G>A p.Met122Ile | |
| NWMN_0144 | | | | | | | | | |
| NWMN_0240 | | | | c.432G>T p.Met144Ile | | | | | |
| NWMN_0309 | c.23A>C p.Asn8Thr | | | | c.23A>C p.Asn8Thr | c.23A>C p.Asn8Thr | | c.23A>C p.Asn8Thr | |
| NWMN_0471 (*tilS*) | | | | | | | | c.760_765dupTTTAAT p.Phe254_Asn255dup | |
| NWMN_0523 (*sdrC*) | | c.2299A>G p.Asn767Asp | | | | | | | |
| NWMN_0564 | | | | | | | c.230_231delCT p.Ser77fs | | |
| NWMN_0954 | | | c.359C>A p.Pro120His | | | | | | |
| NWMN_1152 | | | | | | | | | |
| NWMN_1311 (*lysA*) | | | | | | | | | |
| NWMN_1425 (*recN*) | | | | | | | | | |
| NWMN_1655 | | c.17G>A p.Arg6His | | c.17G>A p.Arg6His | c.17G>A p.Arg6His | c.17G>A p.Arg6His | | c.17G>A p.Arg6His | |
| NWMN_1735 | | | | c.2754C>A p.Phe918Leu | | | | | |
| NWMN_1754 | | | | | | | | c.232T>A p.Leu78Ile | |
| NWMN_2040 (*pdp*) | | c.426G>T p.Leu142Phe | | | | | | | |
| NWMN_2253 | | | | | | | | | c.1412C>T p.Ala471Val |
| NWMN_2388 | | | | c.195G>T p.Gln65His | | | | | |
| POM-HEX $IC_{50}$ (μM) | 8.353 | 31.66 | 11.47 | 6.746 | 4.033 | 5.61 | 5.559 | 3.882 | 2.788 |
| Std. Deviation | 3.673 | 32.94 | 4.009 | 2.199 | 1.276 | 5.394 | 2.96 | 1.153 | 0.538 |

| **Gene (Newman)** | **R19** | **R20** | **R21** | **R22** | **R23** | **R24** | **R25** |
|---|---|---|---|---|---|---|---|
| NWMN_1449 (*gloB*) | | | | | | c.401C>T p.Pro134Leu | |
| NWMN_2528 (*estA, frmB*) | c.40G>C p.Gly14Arg | | c.366G>A p.Met122Ile | | c.356G>A p.Gly119Asp | | c.40G>C p.Gly14Arg |
| NWMN_0144 | | | | | | | |
| NWMN_0240 | | | | | | | |
| NWMN_0309 | c.23A>C p.Asn8Thr | | | | | c.23A>C p.Asn8Thr | c.23A>C p.Asn8Thr |
| NWMN_0471 (*tilS*) | | | | | | | |
| NWMN_0523 (*sdrC*) | | | | | | | |
| NWMN_0564 | | | | | | | |
| NWMN_0954 | | | | | | | |
| NWMN_1152 | | | | | | c.1570G>T p.Asp524Tyr | |
| NWMN_1311 (*lysA*) | | | | | | c.1145C>A p.Ser382Tyr | |
| NWMN_1425 (*recN*) | c.1130T>C p.Leu377Ser | | | | | | |
| NWMN_1655 | c.17G>A p.Arg6His | c.17G>A p.Arg6His | c.17G>A p.Arg6His | c.17G>A p.Arg6His | c.17G>A p.Arg6His | | |
| NWMN_1735 | | | | | | | |
| NWMN_1754 | | | c.232T>A p.Leu78Ile | | | | |
| NWMN_2040 (*pdp*) | | | | | | | |
| NWMN_2253 | | | | | | | |
| NWMN_2388 | | | | | | | |
| POM-HEX $IC_{50}$ (μM) | 5.678 | 3.847 | 3.923 | 4.568 | 3.794 | 8.396 | 3.519 |
| Std. Deviation | 2.073 | 1.061 | 1.087 | 4.194 | 0.903 | 1.049 | 0.112 |

**Supplementary Table 3** Half-maximal inhibitory concentration ($IC_{50}$) values for POM-HEX against transposon mutations in genes identified by whole genome sequencing. Assays performed in biological triplicate with technical duplicates. P-value calculated as a one-way ANOVA with Dunnett correction for multiple comparisons.

| SAUSA300 gene | Newman gene | StrID (NARSA) | Pan gene symbol | POM-HEX $IC_{50}$ (μM) | POM-HEX $IC_{50}$ SD (μM) | Significantly different from WT (JE2)? | Adjusted P-value | Predicted function |
|---|---|---|---|---|---|---|---|---|
| Parental Strain | | JE2 | | 1.54 | 0.40 | | | N/A |
| SAUSA300_1781 | NWMN_1723 | NE64 | *hemY* | 1.43 | 0.15 | ns | 0.9998 | protoporphyrinogen oxidase |
| SAUSA300_0671 | NWMN_0654 | NE364 | | 1.65 | 0.72 | ns | 0.9998 | ABC transporter, ATP-binding protein, MsbA family |
| SAUSA300_1505 | NWMN_1449 | NE377 | *gloB* | 10.6 | 3.2 | **** | <0.0001 | hydroxyacylglutathione hydrolase |
| SAUSA300_1708 | NWMN_1655 | NE386 | *rot* | 1.40 | 0.15 | ns | 0.9997 | staphylococcal accessory regulator Rot |
| SAUSA300_2564 | NWMN_2528 | NE503 | *estA, frmB* | 6.28 | 2.25 | **** | <0.0001 | tributyrin esterase |
| SAUSA300_1452 | NWMN_1410 | NE520 | *proC* | 1.47 | 0.75 | ns | 0.9999 | pyrroline-5-carboxylate reductase |
| SAUSA300_0201 | NWMN_0144 | NE541 | | 1.60 | 0.52 | ns | >0.9999 | peptide ABC transporter, permease protein |
| SAUSA300_1085 | NWMN_1101 | NE874 | | 1.64 | 0.57 | ns | 0.9998 | conserved hypothetical protein |
| SAUSA300_2105 | NWMN_2057 | NE929 | *mtlF* | 1.70 | 0.55 | ns | 0.9997 | PTS system, mannitol specific IIBC component |
| SAUSA300_0778 | NWMN_0762 | NE1051 | | 1.46 | 0.25 | ns | 0.9999 | hypothetical protein |
| SAUSA300_1290 | NWMN_1308 | NE1118 | *dapD* | 1.69 | 0.59 | ns | 0.9997 | tetrahydrodipicolinate acetyltransferase |
| SAUSA300_0414 | NWMN_0407 | NE1127 | *lpl4* | 1.72 | 0.62 | ns | 0.9997 | tandem lipoprotein |
| SAUSA300_0028 | | NE1283 | *tnp* | 2.10 | 0.36 | ns | 0.9986 | putative transposase |

**Supplementary Table 4** Michaelis-Menten parameters for *Sa*GloB. Displayed are the results of three independent biological replicates in technical duplicate.

| | $V_{max}$ (pmol*min$^{-1}$*µg protein$^{-1}$) | | $K_m$ (µM) | | $V_{max}/K_m$ (pmol*min$^{-1}$*mg GloB$^{-1}$µM$^{-1}$) | | $k_{cat}$ ($10^{-3}$ s$^{-1}$) | | $k_{cat}/K_m$ (M$^{-1}$ s$^{-1}$) | | $k_{cat}/k_{uncat}$ ($10^3$) | | $((k_{cat}/K_m)/k_{uncat})$ ($10^9$ M$^{-1}$) | | $((V_{max}/K_m)/k_{uncat})$ ($10^{12}$pmol*mg GloB$^{-1}$µM$^{-1}$) | |
|---|---|---|---|---|---|---|---|---|---|---|---|---|---|---|---|---|
| **Substrate** | **Value** | **SEM** | **Value** | **SEM** | **Value** | **SEM** | **Value** | **SEM** | **Value** | **SEM** | **Value** | **SEM** | **Value** | **SEM** | **Value** | **SEM** |
| 1C | 0.497 | 0.074 | 6.23 | 3.39 | 79.9 | 22 | 0.0744 | 0.0111 | 11.9 | 3.28 | 21.2 | 16.3 | 3.41 | 4.81 | 380 | 536 |
| 2C | 0.292 | 0.051 | 18.3 | 9.0 | 15.9 | 5.6 | 0.0436 | 0.0076 | 2.38 | 0.845 | 17.2 | 7.9 | 0.940 | 0.881 | 105 | 98 |
| 3C | | | >50 | | 10.4 | 1.0 | | | 1.55 | 0.15 | | | 0.256 | 0.061 | 28.5 | 6.8 |
| 6C | 0.444 | 0.130 | 48.0 | 29.5 | 9.24 | 4.40 | 0.0663 | 0.0194 | 1.38 | 0.658 | 12.7 | 33.4 | 0.265 | 1.133 | 29.6 | 126.3 |
| 7C | | | >50 | | 6.43 | 0.31 | | | 0.961 | 0.046 | | | 0.109 | 0.041 | 12.2 | 4.6 |
| 8C | 0.351 | 0.113 | 84.9 | 47.9 | 4.13 | 2.36 | 0.0524 | 0.0169 | 0.618 | 0.353 | 79.1 | 26.3 | 0.932 | 0.549 | 104 | 61 |
| 9C | | | >50 | | 0.28 | 0.01 | | | 0.042 | 0.002 | | | 0.0244 | 0.0020 | 2.72 | 0.22 |
| 10C | | | >50 | | 0.79 | 0.07 | | | 0.118 | 0.011 | | | 0.0752 | 0.0392 | 8.38 | 4.37 |
| 11C | | | >50 | | 0.88 | 0.04 | | | 0.132 | 0.005 | | | 0.261 | 0.006 | 29.1 | 0.7 |
| 1O | 217 | 41.5 | 73.5 | 25.7 | 2960 | 1620 | 32.5 | 6.2 | 442 | 242 | 38.6 | 15.3 | 0.525 | 0.595 | 58.6 | 66.5 |
| 2O | 46.4 | 9.95 | 42.0 | 19.8 | 1103 | 503 | 6.93 | 1.49 | 165 | 75.3 | 19.4 | 11.8 | 0.462 | 0.595 | 51.5 | 66.4 |
| 3O | 4.21 | 0.515 | 8.41 | 3.52 | 500. | 146 | 0.630 | 0.077 | 74.8 | 21.9 | 4.91 | 1.41 | 0.584 | 0.400 | 65.0 | 44.6 |
| 4O | 1.39 | 0.15 | 18.0 | 5.37 | 77.2 | 27.3 | 0.208 | 0.022 | 11.5 | 4.08 | 8.42 | 5.18 | 0.468 | 0.964 | 52.1 | 107.5 |
| 5O | 0.731 | 0.15 | 21.2 | 11.8 | 34.5 | 12.7 | 0.109 | 0.023 | 5.15 | 1.91 | 17.0 | 24.1 | 0.803 | 2.038 | 89.5 | 227.2 |
| 6O | 2.91 | 0.26 | 12.0 | 3.3 | 242 | 76.8 | 0.435 | 0.038 | 36.1 | 11.5 | 5.76 | 1.09 | 0.479 | 0.325 | 53.4 | 36.3 |
| 7O | 23.1 | 4.50 | 20.5 | 10.9 | 1130 | 412 | 3.45 | 0.67 | 168 | 61.7 | 23.6 | 21.2 | 1.15 | 1.95 | 129 | 217 |
| 8O | 1.01 | 0.149 | 15.5 | 6.8 | 64.9 | 22.0 | 0.151 | 0.022 | 9.71 | 3.30 | 6.80 | 1.80 | 0.437 | 0.267 | 48.7 | 29.7 |
| 9O | 2.54 | 0.83 | 55.5 | 36.5 | 45.7 | 22.8 | 0.380 | 0.125 | 6.84 | 3.41 | 14.6 | 16.0 | 0.264 | 0.439 | 29.4 | 48.9 |
| 10O | | | >50 | | 2.96 | 0.22 | | | 0.442 | 0.033 | | | 0.0715 | 0.0346 | 7.97 | 3.85 |
| 11O | | | >50 | | 248 | 21.9 | | | 37.1 | 3.27 | | | 0.0411 | 0.0369 | 4.58 | 4.12 |
| 12O | 1.04 | 0.190 | 36.9 | 15.4 | 28.2 | 12.3 | 0.156 | 0.028 | 4.21 | 1.84 | 1.28 | 0.67 | 0.0348 | 0.0438 | 3.87 | 4.88 |
| 1S | 14.9 | 3.92 | 32.5 | 20.4 | 457 | 192 | 2.22 | 0.59 | 68.4 | 28.8 | 11.4 | 13.6 | 0.351 | 0.666 | 39.2 | 74.3 |
| 2S | | | >50 | | 422 | 39.5 | | | 63.1 | 5.91 | | | 0.0529 | 0.0665 | 5.90 | 7.41 |
| 3S | | | >50 | | 31.0 | 2.34 | | | 4.63 | 0.350 | | | 0.0259 | 0.0088 | 2.89 | 0.98 |
| 4S | | | >50 | | 126 | 15.2 | | | 18.8 | 2.27 | | | 0.278 | 0.057 | 30.9 | 6.4 |
| 5S | 0.18 | 0.049 | 31.7 | 21.1 | 5.60 | 2.34 | 0.0266 | 0.0074 | 0.840 | 0.351 | 4.35 | 2.45 | 0.137 | 0.116 | 15.3 | 12.9 |
| 6S | 4.19 | 1.17 | 65.8 | 34.9 | 63.7 | 33.6 | 0.627 | 0.176 | 9.53 | 5.03 | 12.9 | 7.8 | 0.197 | 0.224 | 21.9 | 25.0 |
| 7S | 0.722 | 0.150 | 49.5 | 21.4 | 14.6 | 7.01 | 0.108 | 0.022 | 2.18 | 1.05 | 7.86 | 3.04 | 0.159 | 0.142 | 17.7 | 15.9 |
| 8S | 1.02 | 0.40 | 79.8 | 56.2 | 12.8 | 7.19 | 0.153 | 0.060 | 1.92 | 1.08 | 9.01 | 7.64 | 0.113 | 0.136 | 12.6 | 15.2 |
| 9S | | | >50 | | 11.1 | 0.86 | | | 1.66 | 0.128 | | | 0.0215 | 0.0613 | 2.40 | 6.84 |
| 11S | | | >50 | | 20.5 | 2.87 | | | 3.07 | 0.429 | | | 0.0662 | 0.0219 | 7.38 | 2.44 |
| 12S | | | >50 | | 1.86 | 0.03 | | | 0.278 | 0.005 | | | 0.0310 | 0.0005 | 3.46 | 0.06 |

**Supplementary Table 5** Michaelis-Menten parameters for *Sa*FrmB. Displayed are the results of three independent biological replicates in technical duplicate.

| | $V_{max}$ (pmol*min$^{-1}$*µg protein$^{-1}$) | | $K_m$ (µM) | | $V_{max}/K_m$ (pmol*min$^{-1}$*mg FrmB$^{-1}$µM$^{-1}$) | | $k_{cat}$ ($10^{-3}$ s$^{-1}$) | | $k_{cat}/K_m$ (M$^{-1}$ s$^{-1}$) | | $k_{cat}/k_{uncat}$ ($10^3$) | | $((k_{cat}/K_m)/k_{uncat})$ ($10^9$ M$^{-1}$) | | $((V_{max}/K_m)/k_{uncat})$ ($10^{12}$ pmol*mg FrmB$^{-1}$µM$^{-1}$) | |
|---|---|---|---|---|---|---|---|---|---|---|---|---|---|---|---|---|
| **Substrate** | **Value** | **SEM** | **Value** | **SEM** | **Value** | **SEM** | **Value** | **SEM** | **Value** | **SEM** | **Value** | **SEM** | **Value** | **SEM** | **Value** | **SEM** |
| 1C | 0.149 | 0.059 | 30.4 | 29.5 | 4.89 | 2.01 | 0.0280 | 0.0112 | 0.921 | 0.379 | 7.99 | 16.5 | 0.263 | 0.557 | 23.3 | 49.3 |
| 2C | 0.0637 | 0.0274 | 26.5 | 28.9 | 2.40 | 0.95 | 0.0120 | 0.0052 | 0.452 | 0.179 | 4.74 | 5.38 | 0.179 | 0.186 | 15.8 | 16.5 |
| 3C | | | >50 | | 10.2 | 0.64 | | | 1.92 | 0.12 | | | 0.316 | 0.050 | 28.0 | 4.4 |
| 6C | | | >50 | | 1.84 | 0.14 | | | 0.347 | 0.026 | | | 0.0666 | 0.0444 | 5.90 | 3.93 |
| 7C | | | >50 | | 1.01 | 0.08 | | | 0.191 | 0.015 | | | 0.0217 | 0.0133 | 1.92 | 1.17 |
| 8C | | | >50 | | 0.314 | 0.012 | | | 0.0591 | 0.0022 | | | 0.0892 | 0.0034 | 7.89 | 0.30 |
| 9C | | | >50 | | 0.160 | 0.009 | | | 0.0301 | 0.0018 | | | 0.0177 | 0.0019 | 1.56 | 0.17 |
| 10C | | | >50 | | 0.233 | 0.029 | | | 0.0438 | 0.0055 | | | 0.0279 | 0.0194 | 2.47 | 1.72 |
| 11C | | | >50 | | 0.682 | 0.062 | | | 0.128 | 0.012 | | | 0.254 | 0.014 | 22.5 | 1.2 |
| 1O | 5.98 | 0.677 | 30.3 | 8.33 | 197 | 81.3 | 1.13 | 0.127 | 37.2 | 15.3 | 1.34 | 0.31 | 0.0442 | 0.0377 | 3.91 | 3.34 |
| 2O | 2.04 | 0.328 | 13.9 | 6.76 | 147 | 48.6 | 0.385 | 0.062 | 27.7 | 9.14 | 1.08 | 0.49 | 0.0777 | 0.0723 | 6.88 | 6.40 |
| 3O | 0.621 | 0.191 | 9.95 | 10.06 | 62.4 | 18.9 | 0.117 | 0.036 | 11.7 | 3.57 | 0.911 | 0.656 | 0.0916 | 0.0653 | 8.11 | 5.78 |
| 4O | 0.183 | 0.038 | 30.0 | 15.2 | 6.12 | 2.51 | 0.0345 | 0.0072 | 1.15 | 0.47 | 1.40 | 1.70 | 0.0467 | 0.1118 | 4.14 | 9.89 |
| 5O | | | >50 | | 1.54 | 0.10 | | | 0.290 | 0.020 | | | 0.0452 | 0.0210 | 4.00 | 1.86 |
| 6O | 0.597 | 0.091 | 9.17 | 4.66 | 65.2 | 19.4 | 0.112 | 0.017 | 12.3 | 3.66 | 1.49 | 0.48 | 0.163 | 0.104 | 14.4 | 9.2 |
| 7O | 2.57 | 0.96 | 107 | 64.8 | 24.0 | 14.8 | 0.485 | 0.180 | 4.52 | 2.78 | 3.31 | 5.69 | 0.0309 | 0.0877 | 2.73 | 7.77 |
| 8O | 0.163 | 0.031 | 18.3 | 9.66 | 8.92 | 3.17 | 0.0307 | 0.0058 | 1.68 | 0.60 | 1.38 | 0.47 | 0.0757 | 0.0482 | 6.70 | 4.27 |
| 9O | | | >50 | | 7.24 | 0.31 | | | 1.36 | 0.06 | | | 0.0525 | 0.0076 | 4.65 | 0.67 |
| 10O | 0.163 | 0.053 | 74.2 | 43.6 | 2.20 | 1.21 | 0.0307 | 0.0099 | 0.414 | 0.227 | 4.97 | 10.26 | 0.0671 | 0.2351 | 5.94 | 20.81 |
| 11O | | | >50 | | 79.1 | 2.7 | | | 14.9 | 0.51 | | | 0.0165 | 0.0058 | 1.46 | 0.51 |
| 12O | | | >50 | | 3.18 | 0.26 | | | 0.598 | 0.048 | | | 0.00493 | 0.00115 | 0.44 | 0.10 |
| 1S | | | >50 | | 12.5 | 0.6 | | | 2.36 | 0.11 | | | 0.0121 | 0.0026 | 1.07 | 0.23 |
| 2S | | | >50 | | 169 | 14 | | | 31.8 | 2.58 | | | 0.0266 | 0.0290 | 2.36 | 2.57 |
| 3S | | | >50 | | 7.79 | 0.19 | | | 1.47 | 0.04 | | | 0.0082 | 0.0009 | 0.728 | 0.081 |
| 4S | 0.505 | 0.147 | 52.7 | 31.3 | 9.58 | 4.70 | 0.0950 | 0.0277 | 1.80 | 0.89 | 1.40 | 0.70 | 0.0266 | 0.0223 | 2.36 | 1.97 |
| 5S | 0.0295 | 0.0030 | 11.1 | 3.61 | 2.66 | 0.83 | 0.00556 | 0.00056 | 0.500 | 0.156 | 0.910 | 0.187 | 0.0818 | 0.0517 | 7.24 | 4.58 |
| 6S | 0.629 | 0.169 | 36.3 | 22.3 | 17.3 | 7.5 | 0.118 | 0.032 | 3.27 | 1.42 | 2.44 | 1.42 | 0.0674 | 0.0633 | 5.97 | 5.61 |
| 7S | | | >50 | | 1.55 | 0.12 | | | 0.292 | 0.023 | | | 0.0213 | 0.0032 | 1.88 | 0.28 |
| 8S | | | >50 | | 2.20 | 0.24 | | | 0.415 | 0.046 | | | 0.0244 | 0.0058 | 2.16 | 0.51 |
| 9S | | | >50 | | 5.38 | 0.48 | | | 1.01 | 0.09 | | | 0.0131 | 0.0435 | 1.16 | 3.85 |
| 11S | | | >50 | | 10.8 | 1.1 | | | 2.04 | 0.21 | | | 0.0440 | 0.0105 | 3.89 | 0.93 |
| 12S | | | >50 | | 1.02 | 0.07 | | | 0.191 | 0.013 | | | 0.0213 | 0.0013 | 1.89 | 0.12 |

**Supplementary Table 6** Summary of crystallographic data collection and refinement statistics.

| Data Collection | *Sa*FrmB | *Sa*GloB (SeMet) |
|---|---|---|
| Space Group | C2 | P 1 21 1 |
| Cell dimensions | a= 128.3Å, b= 80.5Å<br>c=66.7Å, β=113.8° | a= 93.7Å, b= 44.8Å<br>c=105.0Å, β=96.7° |
| Wavelength (Å) | 1.000 | 1.000 |
| Resolution (Å) (highest shell) | 36.8-1.60 (1.63-1.60) | 48 - 1.71 (1.74 - 1.71) |
| Reflections (total/unique) | 145,907 / 78193 | 536,305 / 97,666 |
| Completeness (highest shell) | 96.4% (99.1%) | 96.1 % (88.8 %) |
| <I/σ> (highest shell) | 27.1 (4.1) | 21.3 (1.4) |
| $R_{sym}$ (highest shell) | 6.6% (57.3%) | 10.4% (54.8%) |
| **Refinement** | | |
| $R_{cryst}$ / $R_{free}$ | 0.156 / 0.179 | 0.225 / 0.251 |
| No. of protein atoms | 4164 | 6323 |
| No. of waters | 496 | 431 |
| No. of ligand atoms | 3 | 28 |
| R.m.s.d., bond lengths (Å) | 0.009 | 0.007 |
| R.m.s.d., bond angles (°) | 1.34 | 1.21 |
| Avg. B-factor (Å$^2$): protein, water, ligand | 25.8, 37.6, 15.9 | 39.6, 44.9, 47.5 |
| Stereochemistry: most favored, allowed, disallowed | 98.4, 1.6, 0 % | 96.2, 3.8, 0 % |

**Supplementary Table 7** Michaelis-Menten parameters for human sera. Displayed are the results of three independent biological replicates in technical duplicate.

| | $V_{max}$ (pmol*min$^{-1}$*mg sera$^{-1}$) | | $K_m$ (μM) | | $V_{max}/K_m$ (pmol*min$^{-1}$*mg sera$^{-1}$μM$^{-1}$) | | (($V_{max}/K_m$)/$k_{uncat}$) ($10^{12}$ pmol*mg sera$^{-1}$μM$^{-1}$) | |
|---|---|---|---|---|---|---|---|---|
| **Substrate** | **Value** | **SEM** | **Value** | **SEM** | **Value** | **SEM** | **Value** | **SEM** |
| 1C | 1.06 | 0.15 | 36.2 | 11.5 | 0.0293 | 0.0128 | 0.139 | 0.315 |
| 2C | 0.356 | 0.191 | 71.5 | 70.6 | 0.00498 | 0.00271 | 0.0328 | 0.0471 |
| 3C | | | >50 | | 0.00161 | 0.00022 | 0.00443 | 0.00151 |
| 6C | | | >50 | | 0.00121 | 0.00013 | 0.00389 | 0.00381 |
| 7C | 0.441 | 0.029 | 64.18 | 8.03 | 0.00688 | 0.00355 | 0.0130 | 0.0531 |
| 8C | | | >50 | | 0.000791 | 0.000062 | 0.0199 | 0.0016 |
| 9C | | | >50 | | 0.000581 | 0.000026 | 0.00568 | 0.00045 |
| 10C | | | >50 | | 0.000483 | 0.000042 | 0.00513 | 0.00251 |
| 11C | | | >50 | | 0.000711 | 0.000027 | 0.0234 | 0.0005 |
| 1O | 68.6 | 25.6 | 64.1 | 45.8 | 1.07 | 0.560 | 0.0212 | 0.0230 |
| 2O | 43.8 | 10.53 | 40.4 | 21.5 | 1.09 | 0.489 | 0.0507 | 0.0645 |
| 3O | 11.7 | 2.81 | 22.2 | 14.2 | 0.529 | 0.198 | 0.0687 | 0.0604 |
| 4O | 5.88 | 1.90 | 44.5 | 30.9 | 0.132 | 0.061 | 0.0894 | 0.2420 |
| 5O | 2.25 | 0.48 | 35.4 | 17.4 | 0.0635 | 0.0274 | 0.165 | 0.489 |
| 6O | 11.7 | 3.82 | 54.6 | 35.9 | 0.215 | 0.107 | 0.0475 | 0.0503 |
| 7O | 5.73 | 1.16 | 30.3 | 14.9 | 0.189 | 0.078 | 0.0216 | 0.0411 |
| 8O | 3.93 | 0.71 | 27.7 | 12.4 | 0.142 | 0.057 | 0.107 | 0.077 |
| 9O | 4.35 | 1.18 | 111.7 | 48.9 | 0.0389 | 0.0242 | 0.0250 | 0.0519 |
| 10O | 1.46 | 0.38 | 63.3 | 31.8 | 0.0230 | 0.0121 | 0.0621 | 0.2080 |
| 11O | | | >50 | | 0.122 | 0.008 | 0.00225 | 0.00158 |
| 12O | 4.00 | 0.68 | 42.2 | 15.7 | 0.0948 | 0.0433 | 0.0130 | 0.0171 |
| 1S | 30.5 | 9.90 | 71.8 | 43.0 | 0.425 | 0.230 | 0.0364 | 0.0890 |
| 2S | | | >50 | | 0.315 | 0.025 | 0.00440 | 0.00472 |
| 3S | 8.80 | 2.92 | 74.3 | 45.0 | 0.118 | 0.065 | 0.0111 | 0.0273 |
| 4S | 3.81 | 0.39 | 27.9 | 7.08 | 0.137 | 0.055 | 0.0336 | 0.0230 |
| 5S | 0.690 | 0.246 | 31.0 | 26.7 | 0.0222 | 0.0092 | 0.0606 | 0.0510 |
| 6S | | | >50 | | 0.0257 | 0.0025 | 0.00886 | 0.00182 |
| 7S | 1.60 | 0.43 | 30.2 | 19.7 | 0.0528 | 0.0216 | 0.0641 | 0.0490 |
| 8S | | | >50 | | 0.0140 | 0.0007 | 0.0138 | 0.0015 |
| 9S | | | >50 | | 0.0247 | 0.0021 | 0.00532 | 0.01651 |
| 11S | | | >50 | | 0.0190 | 0.0017 | 0.00682 | 0.00149 |
| 12S | | | >50 | | 0.00504 | 0.00036 | 0.00937 | 0.00062 |

**Supplementary Table 8** Michaelis-Menten parameters for mouse sera. Displayed are the results of three independent biological replicates in technical duplicate.

| | $V_{max}$ (pmol*min$^{-1}$*mg sera$^{-1}$) | | $K_m$ (μM) | | $V_{max}/K_m$ (pmol*min$^{-1}$*mg sera$^{-1}$μM$^{-1}$) | | $((V_{max}/K_m)/k_{uncat})$ ($10^{12}$ pmol*mg sera$^{-1}$μM$^{-1}$) | |
|---|---|---|---|---|---|---|---|---|
| **Substrate** | **Value** | **SEM** | **Value** | **SEM** | **Value** | **SEM** | **Value** | **SEM** |
| 1C | | | >50 | | 4.56 | 0.51 | 21.7 | 12.4 |
| 2C | | | >50 | | 1.62 | 0.33 | 10.7 | 5.7 |
| 3C | | | >50 | | 0.52 | 0.02 | 1.43 | 0.13 |
| 6C | 17.6 | 9.4 | 53.5 | 58.1 | 0.33 | 0.16 | 1.05 | 4.65 |
| 7C | | | >50 | | 2.04 | 0.27 | 3.86 | 4.04 |
| 8C | | | >50 | | 1.35 | 0.13 | 34.0 | 3.3 |
| 9C | 3.18 | 0.66 | 12.3 | 8.0 | 0.26 | 0.08 | 2.53 | 1.46 |
| 10C | | | >50 | | 0.93 | 0.08 | 9.87 | 4.74 |
| 11C | | | >50 | | 0.28 | 0.01 | 9.34 | 0.18 |
| 1O | | | >50 | | 14.01 | 0.83 | 0.28 | 0.03 |
| 2O | | | >50 | | 15.90 | 1.12 | 0.74 | 0.15 |
| 3O | | | >50 | | 57.0 | 43.2 | 7.41 | 13.17 |
| 4O | | | >50 | | 6.76 | 0.21 | 4.57 | 0.82 |
| 5O | | | >50 | | 20.3 | 12.0 | 52.6 | 214.7 |
| 6O | 437 | 135 | 11.5 | 8.65 | 38.1 | 15.6 | 8.42 | 7.39 |
| 7O | 929 | 103 | 25.9 | 7.30 | 35.8 | 14.1 | 4.08 | 7.40 |
| 8O | | | >50 | | 15.7 | 0.4 | 11.8 | 0.5 |
| 9O | | | >50 | | 6.69 | 0.17 | 4.30 | 0.36 |
| 10O | | | >50 | | 2.93 | 0.14 | 7.91 | 2.36 |
| 11O | | | >50 | | 3.24 | 0.16 | 0.06 | 0.03 |
| 12O | | | >50 | | 11.9 | 0.4 | 1.64 | 0.15 |
| 1S | | | >50 | | 14.2 | 1.4 | 1.22 | 0.54 |
| 2S | | | >50 | | 5.07 | 0.21 | 0.07 | 0.04 |
| 3S | | | >50 | | 9.47 | 0.76 | 0.88 | 0.32 |
| 4S | | | >50 | | 24.2 | 2.6 | 5.96 | 1.08 |
| 5S | | | >50 | | 1.93 | 0.13 | 5.27 | 0.71 |
| 6S | | | >50 | | 12.1 | 0.6 | 4.16 | 0.41 |
| 7S | | | >50 | | 11.9 | 1.1 | 14.4 | 2.6 |
| 8S | | | >50 | | 5.17 | 0.43 | 5.07 | 0.91 |
| 9S | | | >50 | | 1.29 | 0.11 | 0.28 | 0.89 |
| 11S | | | >50 | | 1.97 | 0.10 | 0.71 | 0.09 |
| 12S | | | >50 | | 5.74 | 0.53 | 10.7 | 0.9 |

**Supplementary Table 9** Primers used during this study.

| No | Name | Sequence | Use |
|---|---|---|---|
| 1 | NWMN_0144_F | TTTTCCTGATCCTGATTCAC | Sanger Sequencing |
| 2 | NWMN_0144_R | ATGATGCTTCCATGTTTGTT | Sanger Sequencing |
| 3 | NWMN_0306_F | AATACACCGGGTAACACAAC | Sanger Sequencing |
| 4 | NWMN_0306_R | CGTTTTGTTGAGCTAATTCC | Sanger Sequencing |
| 5 | NWMN_0309_F | ACCATGCTTAAAGGGATTTT | Sanger Sequencing |
| 6 | NWMN_0309_R | TGTCACCTAAGTCAACACCA | Sanger Sequencing |
| 7 | NWMN_0407 (*lpl4nm*) _F | CCGTTGGAGATAGGAAGTTA | Sanger Sequencing |
| 8 | NWMN_0407 (*lpl4nm*) _R | TTTGTGCTTCTTTTGAACCT | Sanger Sequencing |
| 9 | NWMN_0654_F | GAAAATGGAAGACTGATTGC | Sanger Sequencing |
| 10 | NWMN_0654_R | TAATGCATCTGACAAAGTCG | Sanger Sequencing |
| 11 | NWMN_0762_F | GGTGAAGTTTTGGACGATAA | Sanger Sequencing |
| 12 | NWMN_0762_R | TTTTCATCTGTCCGACTTTT | Sanger Sequencing |
| 13 | NWMN_1101_F | TCCACCTATTGGAATTATCG | Sanger Sequencing |
| 14 | NWMN_1101_R | AGACGTTCAATTTCAGTGCT | Sanger Sequencing |
| 15 | NWMN_1192 (*pgsA*) _F | TGGGACGAAGTAATTACAGT T | Sanger Sequencing |
| 16 | NWMN_1192 (*pgsA*) _R | ATATCCCCCTTGTATCGTTT | Sanger Sequencing |
| 17 | NWMN_1308 (*dapD*) _F | TCTATTCGTGGAGGTACGAT | Sanger Sequencing |
| 18 | NWMN_1308 (*dapD*) _R | ATCGTATGTGAGCCATTACC | Sanger Sequencing |
| 19 | NWMN_1410_F | CGATAAACCTAAACCACTCG | Sanger Sequencing |
| 20 | NWMN_1410_R | ATAAACAATGCTTGCCAAAT | Sanger Sequencing |
| 21 | NWMN_1505_F | TGAAGGTGAATTAAGCGATG | Sanger Sequencing |
| 22 | NWMN_1505_R | TGCTATTCCCAATTTGTTCA | Sanger Sequencing |
| 23 | NWMN_1655_F | GAATTGTTGCAATTTAATGGT | Sanger Sequencing |
| 24 | NWMN_1655_R | AACGTAATCATGCTCCATTC | Sanger Sequencing |
| 25 | NWMN_1679_F | CCATGGGAAAAATTAGACAA | Sanger Sequencing |
| 26 | NWMN_1679_R | AAATATCGCCTCACCTTTTT | Sanger Sequencing |
| 27 | NWMN_1723 (*hemY*) _F | GCCGAATACACATCCATTAT | Sanger Sequencing |
| 28 | NWMN_1723 (*hemY*) _R | AACCTTTGTCTCTGCTTCAA | Sanger Sequencing |
| 29 | NWMN_1851 (*nadC*) _F | AGCCATTTTAGCACCATAAA | Sanger Sequencing |
| 30 | NWMN_1851 (*nadC*)_R | TAGAATCCTGTCCTCCTGAA | Sanger Sequencing |
| 31 | NWMN_2057 (*mtlF*)_F | TGTACAACGGTGTTGTTTTG | Sanger Sequencing |
| 32 | NWMN_2057 (*mtlF*)_R | CGGTGAATAGTACGAGAGGA | Sanger Sequencing |
| 33 | NWMN_2528_F | ACTGATGCTTTACCAGAAAC | Sanger Sequencing |
| 34 | NWMN_2528_R | TCAGCGGTAGTAATAAAGGT | Sanger Sequencing |

**Supplementary Table 10** Accession numbers for the isolates used in WhatsGNU analysis.

| Isolate | Bioproject | Biosample | WGS | SRA |
|---|---|---|---|---|
| AD_3_179 | PRJNA512846 | SAMN10689346 | VYMI00000000 | SRR8389007 |
| AD_11_548 | PRJNA512846 | SAMN10689354 | VYMN00000000 | SRR8389002 |
| AD_14_565 | PRJNA512846 | SAMN10689355 | VYMO00000000 | SRR8389003 |
| AD_16_660 | PRJNA512846 | SAMN10689358 | SJAX00000000 | SRR8389044 |
| AD_61_868 | PRJNA512846 | SAMN10689401 | VYNS00000000 | SRR11016776 |
| AD_85_830 | PRJNA512846 | SAMN10689428 | VYOK00000000 | SRR8389056 |
| AD_96_471 | PRJNA512846 | SAMN10689447 | VYOW00000000 | SRR8389016 |
| AD_103_347 | PRJNA512846 | SAMN10689453 | VYPC00000000 | SRR8389035 |
| AD_113_782 | PRJNA512846 | SAMN10689463 | VYPL00000000 | SRR8389099 |
| SSTI_227_44 | PRJNA563582 | SAMN12642230 | VUGB00000000 | SRR11016228 |
| SSTI_228_42 | PRJNA563582 | SAMN12642226 | VUGF00000000 | SRR11016232 |
| SSTI_231_2 | PRJNA563582 | SAMN12642218 | VUGN00000000 | SRR11016241 |
| SSTI_233_51 | PRJNA563582 | SAMN12642215 | VUGQ00000000 | SRR11016244 |
| SSTI_235 | PRJNA563582 | SAMN12642210 | VUGV00000000 | SRR11016250 |
| SSTI_241_9 | PRJNA563582 | SAMN12642203 | VUHC00000000 | SRR11016198 |
| SSTI_247_75 | PRJNA563582 | SAMN12642193 | VUHM00000000 | SRR11016209 |
| SSTI_258_57 | PRJNA563582 | SAMN12642177 | VUIB00000000 | SRR11016226 |
| SSTI_290 | PRJNA563582 | SAMN12642174 | VUIE00000000 | SRR11016283 |
